# fastder: fast, annotation-agnostic detection of expressed regions from RNA-seq coverage and splicing data

**DOI:** 10.64898/2026.06.21.733617

**Authors:** Martina Lehmann, Tom Kitak, Izaskun Mallona

**Affiliations:** Department of Molecular Life Sciences, University of Zurich, Zurich, Switzerland; SIB Swiss Institute of Bioinformatics, Zurich, Switzerland

## Abstract

**Background.:** Features from RNA-seq experiments are usually counted using a curated annotation (GENCODE, Ensembl, etc) as reference. This approach constrains RNA-seq counts to what the annotation considers a valid gene, transcript or exon, and misses transcription outside them. Moreover, reference annotations are known to be incomplete, and inadequate in some experiments altering splicing, and some diseases. To circumvent these constraints, annotation-free detection of expressed regions aims to detect the actual regions being expressed in a sample or samples. For large scale usage, the `recount3` resource holds uniformly processed coverage tracks and splice junctions for over 8,000 human and over 10,000 mouse RNA-seq studies, with several hundred thousand samples in total. This opens an opportunity for scalable tools to call expressed regions that are annotation free.

**Results.:** We present `fastder`, a C++ tool to detect expressed regions directly from `recount3` coverage and splice junctions files, with no read alignment steps. It also can run on local raw reads, after aligning them with `STAR`, making it suitable to analyze species beyond human and mouse. `fastder` calls expressed regions by bump hunting expression coverage files, and stitches them into spliced multi-exon structures using the splice junction data. On simulated RNA-seq with unannotated expression features, `fastder` calls exons at a base-level accuracy on par with `derfinder` and golden standard for the task. Exon precision improves with sequencing depth. We compare `fastder`’s performance against a coverage-only baseline, `derfinder` and `groHMM`. `fastder` extends the functionality of those by assigning strands to the expressed regions, and by also producing multi-exon regions. It runs about 15 to 20 times faster than `derfinder` and `groHMM`, within bounded memory. We showcase its use with two `recount3`-derived examples, including the recovery of cryptic exons from a TDP-43 knockdown experiment; and the GTEx data clustering based on the topology of the expressed regions alone, without taking the expression levels into account.

**Conclusions.:** `fastder` makes annotation-agnostic detection of expressed-regions fast enough to run at `recount3` scale. As limitation, it detects expressed regions and the junctions between them, not different isoforms from the same gene.

## Introduction

RNA sequencing (RNA-seq) data analysis aims to quantify gene expression levels and link them to underlying biological phenotypes [1]. A common approach is to align short reads to the genome and relate them to existing gene annotations such as GENCODE [2] or RefSeq [3]. Existing gene annotations remain incomplete and fail to capture the full complexity of the transcriptome [4]. Unannotated transcription is most often described in the human brain, where is a limiting factor in the mechanistic understanding of neurological diseases [4]: up to 41% of transcription in the human frontal cortex remains unannotated [5]. The CHESS database, built from about 10’000 GTEx RNA-seq experiments, added thousands of novel transcripts absent from RefSeq or GENCODE [6, 7]. The number of protein-coding genes and, especially of transcripts, still varies widely across annotation databases [7].

Existing gene annotations are also often incorrect. About a third of the reported gene rearrangements in mammalian mitochondrial genome annotations in the NCBI database are reported to be errors [8]. Relying on existing annotations to infer gene expression also hinders the discovery of new transcripts [5]. Building a complete and correct annotation of the human genome is an ongoing process, which is why annotation-independent approaches in RNA-seq analysis stay useful. This advantage is larger for nonmodel species with incomplete annotations.

Some tools already detect transcription without an annotation. `derfinder` uses read-count bump hunting combined with a statistical model to identify differentially expressed regions [9]. `TAR-scRNA-seq` builds on a Hidden Markov model to segment the genome into transcribed and untranscribed regions from RNA-seq coverage[10]. Both are moderately heavy in computation, and neither were designed to read precomputed coverage summaries at large scale, such as those from `recount3`. `recount3` removes the alignment and coverage steps for public data [11], as it provides uniformly processed RNA-seq data for over 8,000 human and over 10,000 mouse studies. Each study has thousands of per-sample coverage bigWig files and one set of per-study splice junction files. Generating annotation-independent RNA-seq results from this resource at scale needs a caller that reads coverage and junctions directly and stays within bounded memory genome-wide.

We present `fastder`, a fast, annotation-agnostic splice-junction-aware C++ tool for the detection of expressed regions. `fastder` takes genome-wide coverage and splice junction files, performs bump hunting on the normalized base-resolution coverage to call expressed regions (ERs), and uses the junctions to chain together expressed regions. It does not reconstruct full isoforms: teasing apart co-expressed isoforms from summed coverage is beyond what coverage and junction counts support. Hence, we compare its performance to similarly aimed coverage-only tools.

## Implementation

`fastder` is written in C++ (C++20). It takes the coverage and splice-junction files produced by `recount3`, or equivalent files from a `Monorail`-style pipeline [11] (named `recount3`-like, also provided by custom helpers with `fastder`), and writes a GTF file of expressed regions, including individual exons and their stitched transcripts (Fig. 1A). The code is organized in three stages, one per stage of the algorithm: parsing, averaging with region detection, and splice-junction integration (Fig. S1B). Performance comparisons are provided in a companion repository against splicing scenarios with known ground truth (Fig. 1B).

**Figure 1:**
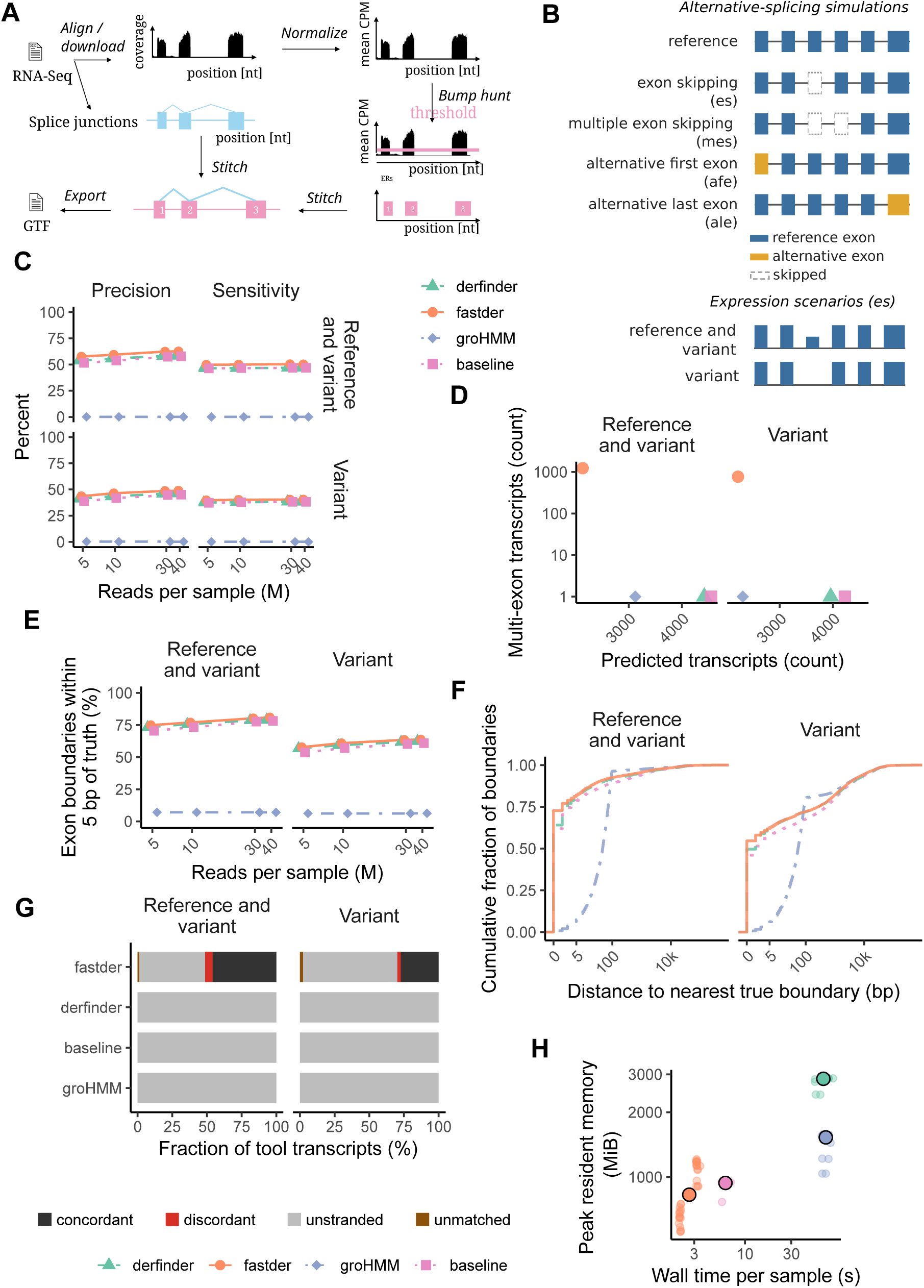
`fastder` architecture and performance. **(A)** The `fastder` pipeline, from `recount3` input files to the GTF of expressed regions. **(B)** Simulation design for performance evaluation. **(C)** Exon-level sensitivity and precision across the sequencing-depth sweep (5, 10, 30 and 40 million reads per sample). **(D)** Multi-exon transcripts called. **(E)** Boundary precision against depth. **(F)** Cumulative distribution of the distance from each predicted boundary to the n3earest reference boundary. **(G)** Strand concordance, pooled over the alternative-splicing samples. **(H)** Wall time and peak resident memory during simulations.

### Input data and recount3 integration

`fastder` consumes four file types from the `recount3` or `recount3`-like workflow. Coverage is required as per-sample bigWig files, a binary format storing continuous RNA-seq coverage along the genome. Splice junctions (as generated by `STAR`, including canonical but also novel junctions detected during the alignment) are supplied as exon-exon junction counts in Matrix Market format, including an MM file with per-sample junction counts and an RR file with junction coordinates, strand, splice motif and annotation flag (e.g., whether the junction is novel or not). A metadata CSV maps the three sample identifiers that `recount3` uses. The file formats are described in Supplementary Section Coverage can be read directly from bigWig through libBigWig, or converted to bedGraph first with `bigWigToBedGraph` using Kent’s utils [12].

### Coverage representation

Coverage is held in memory as a sparse list of intervals: on human chr21 it requires a resident set in the low hundreds of megabytes, against few gigabytes for a dense representation at full hg38. The memory cost scales with the number of distinct coverage intervals rather than with genome length, keeping `fastder` runs within bounded memory.

### Expressed region detection

After parsing, `fastder` computes a per-nucleotide mean coverage vector for each chromosome, with one thread per chromosome in parallel runs. The number of parallel tasks (chromosomes) is specified by the user. It then applies bump hunting to the mean coverage (Fig. 1A): uninterrupted genomic intervals with normalized coverage at or above `min_coverage` and a length of at least `min_length` are called an expressed region. Coverage is normalized to counts per million (CPM) using the per-sample library size, enabling an interpretable expression threshold across samples and experiments.

### Splice-junction stitching

`fastder` then stitches expressed regions across splice junctions. Two regions are joined into a stitched expressed region (e.g., a spliced transcript) when a junction in the input connects the end of a region to the start of the next, within a coordinate tolerance set by `position_tolerance` (Fig. S1C). The stitching is annotation-agnostic by construction: it uses every junction in the `recount3` MM/RR matrix, annotated and novel alike, without filtering for or weighting either by their annotation. Junctions are weighted by their read coverage. The rationale behind not stitching based on novel junctions alone is to keep a low false-discovery rate and reduce the influence of the annotation used during read alignment.

When a stitched chain crosses a junction, the exon edges on either side are set to the junction donor and acceptor coordinates, rather than to the coverage-inferred edges of the expressed region. A stitched exon boundary hence sits on the splice site, not on the coverage decay alone. This behaviour is intended to capture splicing events such as cryptic exons. The outer ends of an expressed regions, and both ends of a single-exon regions, define their boundaries based on coverage drops. Stitching is strand-aware, so stitched expressed regions inherit the strand of the junction(s) used for stitching. Unstitched regions stay unstranded.

### Concurrency and parsing

File parsing, mean coverage computation and bump hunting run concurrently. `fastder` uses `std::thread` so each chromosome or scaffold is processed by exactly one thread and idle threads pick up the next unprocessed chromosome. The parsing stage is a large part of the runtime, because the input files can be large: an MM file provided by `recount3` can hold ca. 700 million lines. Integer columns are parsed with `std::from_chars`, and the mean coverage vector is preallocated at its known size. The parsing and concurrency handling are described in Supp. Methods.

### Coordinates convention

`fastder` works internally in 0-based, half-open coordinates, the convention for bigWig and bedGraph. RR splice-junction coordinates, following `recount3`’s representation, are 1-based and closed. Output GTFs are 1-based and closed, as per standard. Conversions are run on read or write.

### Benchmarking

Performance data were collected with the benchmark directive from Snakemake on runs on a Linux server with an AMD EPYC 7742 64-core processor and 503 GiB of RAM.

## Results

### Performance on simulated data with known expressed regions

We benchmarked `fastder` against three methods: `derfinder` [9], `groHMM` [13] (the expressed region finder behind `TAR-scRNA-seq` [10]), and a baseline segmentation tool based on coverage-only, named the `megadepth` baseline, that emits regions above a expression threshold with no junction stitching. All four tools read the same coverage bigWig files and use the same CPM normalization. The `megadepth` baseline provides a yardstick to quantify the contribution of splice junction stitching (present only in `fastder`), whereas `groHMM` and `derfinder` are well performing standards in the field. To generate data with known truths, we simulated raw RNA-seq reads using `ASimulatoR` [14] under specific splicing anomaly scenarios (Fig. 1B), restricted to chr21, and aligned the reads with the `Monorail` workflow [11], which `recount3` also uses. Performance metrics were computed with `gffcompare` [15], including metrics for matching exon boundaries; and also with fuzzy metrics, including the best Jaccard overlap, distance to the nearest boundary, locus recall, and strand agreement. The evaluation pipeline is part of a Snakemake workflow accompanying `fastder`. The workflow handles reproducible software environments using conda and singularity. Default parameters across tools included an expression threshold of `min_coverage` 0.05 CPM to call regions, `min_length` 10 bp and `position_tolerance` 5 bp for `fastder`; the matching specifications for coverage threshold for `derfinder` and the `megadepth` baseline; and `groHMM`’s best-Jaccard grid point on the simulation, since its LtProbB/UTS parameters do not encode coverage thresholds directly.

We first simulated reads with known splicing anomalies at a depth of 10 million reads per sample. `fastder` called expressed regions at 94.2% base precision and 60.0% base sensitivity, performing similarly to `derfinder` (96.3% and 61.7%) and the `megadepth` baseline (96.3% and 61.7%). At the exon level, `derfinder` (59.2%) and the baseline (58.8%) performed better than `fastder` (56.2%). At the transcript and locus level the metrics were very low for `fastder` (1.3% transcript precision, 5.8% locus sensitivity) and zero for `derfinder`, the baseline and `groHMM`, which emit single-exon (not multiexon) features (Table S1). `groHMM` showed the highest base sensitivity (83.7%) but near-zero exon precision (0.2%), because its HMM scans coverage files binned in 50 nt-long tiles that are unlikely to match exon boundaries. As expected, each tool showed different yet complementary strengths and limitations.

We next evaluated the base-level accuracy across a range of library sizes to evaluate the impact of deeper sequencing/expression level, including 5, 10, 30 and 40 million reads per sample. Results were largely unchanged across that range, with ca. 94% base precision and 60% base sensitivity for `fastder`, ca. 96% and 62% for `derfinder` and the baseline. Exon precision improved with depth, from 53.7% at 5 million reads to 62.1% at 40 million for `fastder`, with a performance advantage over `derfinder` and the `megadepth` baseline at 30 and 40 million reads (Table S2, Fig. 1C,E). This is consistent with low coverage being a good enough indicator to place exon boundaries, whereas a higher coverage is needed to sequence enough spliced reads, which are key to stitch exons across junctions. In terms of transcript, intron-chain and locus levels metrics as calculated by `gffcompare`, `derfinder` and the baseline scored zero by construction, because they emit single-exon regions, and `groHMM` scored near zero because of the poor precision in finding exon boundaries (Figs. S2, S3, S4). Most missed features were shared by all four tools (Fig. S5). The performance gain of `fastder` with deeper library sizes arose from stitching single exons into multi-exon structures using spliced reads, yet most `fastder` regions remained monoexonic (Fig. S6), with a tail of multiexon expressed regions (Fig. 1D) that span a range of exon counts, exonic lengths and mean coverages (Figs. S7, S8, S9). Visually, the called expressed regions resembled the true simulated loci as genome browser tracks (Fig. S10).

**Figure 2:**
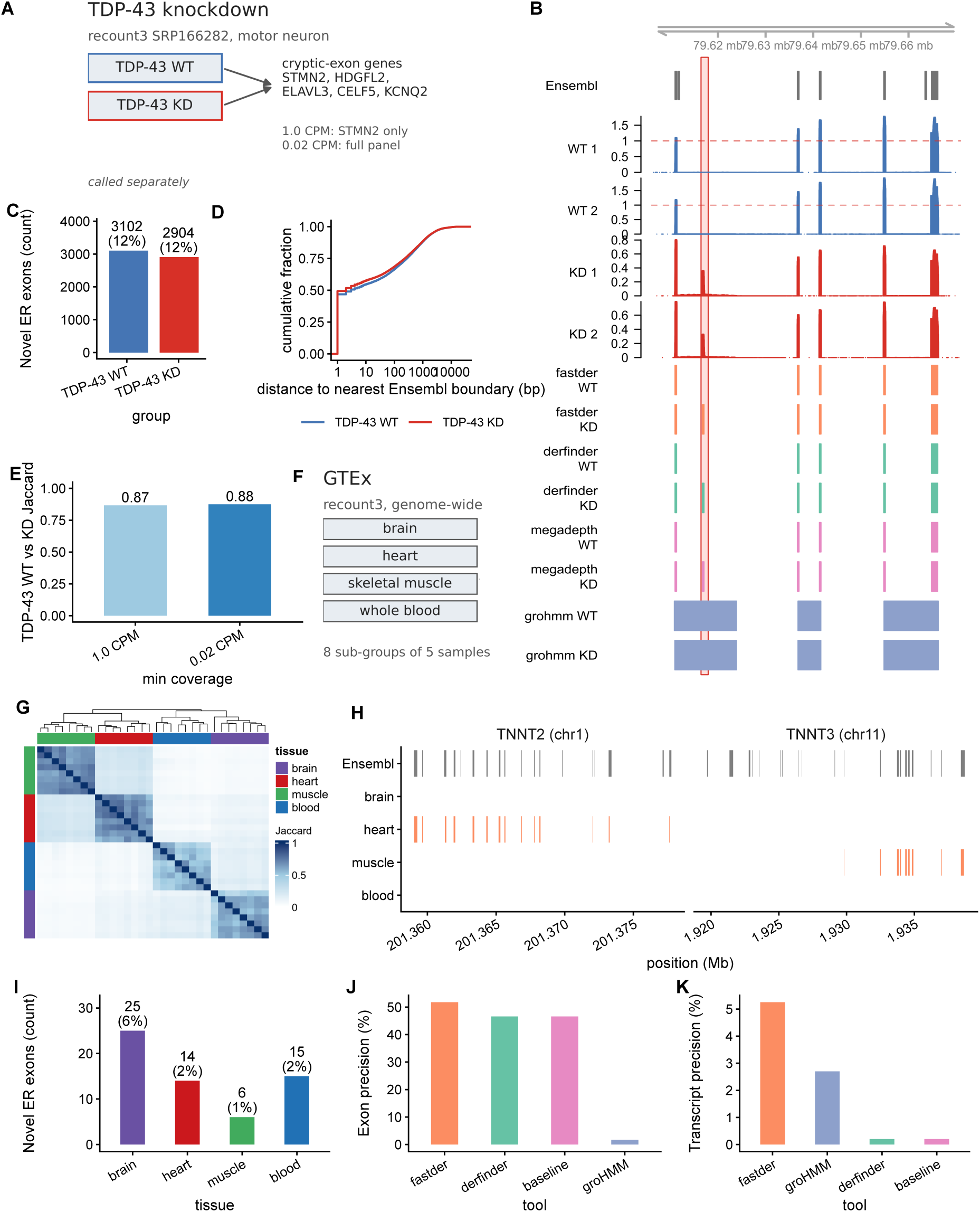
`fastder` applications. **(A)** Experimental design of the TDP-43 knockdown (`recount3` SRP166282) example. **(B)** The STMN2 cryptic exon recovered by `fastder`: tracks show the Ensembl gene models, per-sample coverage for knockdown and control, and each tool’s called regions; the shaded box marks the cryptic exon, absent from Ensembl and called only in the knockdown. **(C)** Novel expressed-region exons per condition (not overlapping Ensembl exons). **(D)** Cumulative distribution of the distance from each `fastder` boundary to the nearest Ensembl boundary, knockdown against control. **(E)** Exonic Jaccard between the knockdown and control at the 1 CPM and 0.02 CPM thresholds. **(F)** Experimental design of the GTEx across four tissues example. **(G)** Structural concordance of the 32 per-sub-group catalogs by exonic Jaccard overlap; catalogs cluster by tissue, with brain the most distinct block. **(H)** Troponin marker *loci* : TNNT2 on chr1 in heart and TNNT3 on chr11 in skeletal muscle. **(I)** Novel expressed-region exons per tissue (not overlapping Ensembl exons). **(J**8) Exon precision and **(K)** transcript precision against Ensembl on the chr19 cross-tool run.

`gffcompare`’s metrics are traditionally used to describe the quality of assembled transcriptomes against a reference annotation, which deviates from our benchmarking task. To extend the evaluation, we devised fuzzy matching metrics (see Suppl. Methods), which do not require exact regional boundaries as `gffcompare` does. The median exonic Jaccard overlap of a called region against the matching true (simulated) transcript was 0.371 for `fastder` and 0.288 for both `derfinder` and the `megadepth` baselines, and 0.38 for `groHMM` (Fig. S11). As with `gffcompare`-derived metrics, the main performance difference was precision and accuracy calling boundaries. `fastder` put 71.5% of its boundaries within 5 bp of a true (simulated) splice site (Fig. S12), against 75.6% for `derfinder` and 75.0% for the baseline, while `groHMM` placed only 7.4% within 5 bp, with a median boundary offset of 46 bp largely related to the coverage binning resolution, set to 50 nt (Figs. 1F and S13). The boundary-distance tolerance had little impact: sweeping `fastder`’s `position` `tolerance` (and `derfinder`’s `maxregiongap`) from 0 to 20 bp did not change the median exonic Jaccard overlap against the simulated truth (Fig. S14).

In summary, `fastder` matched `groHMM` performance in region overlap with the true expressed regions, while improving in the boundary placement. In terms of locus recall at 50% exonic overlap, `fastder`, `derfinder` and the coverage-only baseline scored ca. 42%, behind `groHMM`’s 72.6% recall, as expected from its coarse, high-coverage segments (Fig. S15). `fastder` assigned a strand to 38.8% of its calls, and 94% of those agreed with the reference strand; no other tools called stranded regions (Fig. 1G).

In terms of computing profiling on the AsimulatoR benchmark at 10M reads, with all four tools reading the same coverage files, `fastder` ran in about 2 to 5 seconds per sample (2.2 s at 5 million reads, 4.6 s at 40 million), against 61 to 72 s for `derfinder` and 68 to 72 s for `groHMM`, about 15 to 20 times slower than `fastder`. The `megadepth` baseline ran in 6 to 9 s, a few times slower than `fastder` (Fig. 1H). Peak resident memory followed the same trend (Table S3).

### Performance on simulated data with known splicing aberrations

The `ASimulatoR` framework simulates five alternative-splicing event classes with their abundances; as a result, it generates fastq reads amenable to the `STAR` aligning workflow bundled with `fastder`. We simulated four samples with one event class each: exon skipping (es), multiple exon skipping (mes), alternative first exon (afe) and alternative last exon (ale); plus a mixed sample that draws events from all classes with equal probability. Each class was simulated under two regimes: both the reference and spliced variant being expressed (named template-and-variant), or without the reference allele (variant-only) (Fig. 1B).

`fastder` detected terminal-exon events with good precision. In the template-and-variant scenario, where both the canonical and the altered isoforms are expressed, and measured according to exonic Jaccard’s overlap, exon precision was highest for alternative first (57.7%) and alternative last exons (57.3%), followed by single exon skipping events (56.2%), being the multiple exon skipping the lowest in performance (53.1%). An alternative terminal exon produces a clean coverage block flanked by one splice junction and one transcription start or end, so both boundaries are easier to call. Multiple exon skipping, on the contrary, leaves several short exons and close junctions in a small window, so missed or shifted boundaries impacts the scoring and increases the difficulty of the task. The variant-only scenario, where the reference isoform is no longer expressed, produced similar results (Table S4): exon skipping and terminal-exon events stayed near 50% exon precision, while multiple exon skipping fell to 32.0%. Multiple exon skipping is the hardest case to call expressed regions, and the terminal-exon events are the easiest. The relative task difficulty was maintained across the tested `fastder` parameters (Fig. S16).

### TDP-43-triggered cryptic exons detection

Loss of TDP-43 is known to trigger the expression of cryptic exons [16]. We ran `fastder` on a TDP-43 knockdown in motor neurons and its matched control from SRP166282 (GEO GSE121569) [17] as processed by `recount3` (e.g., without alignment) (Fig. 2A). We explored the recovery of a curated list of genes with reported cryptic exons in TDP-43 knockdowns. These cryptic exons, albeit recurrent, remain unannotated in the standard annotations. `fastder` recovered the STMN2 cryptic exon (Fig. 2B), as well as HDGFL2 (Fig. S17), ELAVL3 (Fig. S18) and KCNQ2 (Fig. S19). While STMN2 cryptic exon was callable based solely on the high expression above the intronic noise signal, the rest of the panel relied on the parsing of spliced junctions for detection. `fastder`’s performance in recovering the cryptic exons was comparable or better than `derfinder`, `groHMM` and the `megadepth` baseline.

The TDP-43 knockdown and control runs were similar, each with about 3,000 novel expressed-region exons (ca. 12% of the catalog, none overlapping an Ensembl exon; Fig. 2C) and nearly identical distance from each `fastder` boundary to the nearest Ensembl boundary (Fig. 2D). The exonic Jaccard between the knockdown and control was 0.87 and 0.88 at a 1 CPM and 0.02 CPM calling thresholds, respectively (Fig. 2E). Hence, the TDP-43 knockdown effect in `fastder`’s reference-free expression regions was minimal, even if known cryptic exon *loci* were recapitulated. In terms of computing performance, `fastder` ran on each sample group in about 4.4 s with ca. 1.3 GB peak memory, a smaller fingerprint than `derfinder`, `groHMM` and the `megadepth` baseline (Table S3).

### Structural similarities across GTEx tissues

To stress test `fastder` performance on larger datasets, we next evaluated GTEx samples [18] comprising brain, heart, muscle and blood tissues. Each tissue group was divided into eight independent subgroups of five `recount3` samples each. Expressed regions were called per subgroup (32 catalogs in total; Fig. 2F). `fastder` detected catalog-specific expressed regions. To evaluate whether these expressed regions recapitulated the tissue of origin, we clustered the catalogs using the pairwise exonic Jaccard similarity of their exons, while neglecting their expression levels. We found that sub-groups of samples from the same tissue grouped together, with brain being the most distinctive block, as expected [4, 5] (Fig. 2G). Hence, `fastder` recapitulated expressed regions shapes that, regardless of their expression level, are characteristic of their tissue of origin. The same tissue separation could also be confirmed by browsing a curated list of marker loci, for instance the cardiac troponin TNNT2 and the skeletal troponin TNNT3 (Fig. 2H), only detectable in heart and muscle, respectively.

We also compared `fastder` to the other tools, while subsetting the chromosomes to chr19 for the sake of speed. As evaluation metric we calculated deviations from each tool’s expressed regions against the Ensembl annotation, using the same 32 catalogs as before. We reasoned that closeness to the curated Ensembl annotation is a good proxy for recovering ‘true’ regions, even if non-canonical expression is expected in brain. All tissues showed some degree of novel expressed regions (not cataloged in the original Ensembl annotation), being brain the most dissimilar (Fig. 2I). `fastder` reached 51.8% exon precision against 46.6% for `derfinder` and the baseline, and 5.2% transcript precision against 0.2% (Fig. 2J,K), consistent with the boundary-snapping and multi-exon behaviour seen on the simulated data. `groHMM` showed a 1.7% exon- and 33.9% base precision, respectively, consistent with a coverage segmentation using tiling data (Table S5). The `gffcompare` sensitivity was only ca. 2% for every tool, as expected due to unfairly comparing the restrictive expressed genes in a tissue over a given expression threshold to the whole Ensembl catalog. On this run `fastder` regions spanned a range of exon lengths (Fig. S20) and emitted mainly unstranded regions (Fig. S21).

In terms of performance on the chr19 subset across tools, `fastder` processed one sample sub-group in a. 4.5 s with peak resident set near 1.5 GB, against 57.6 s and ca. 3.4 GB for `derfinder`, 75.8 s and ca. 1.2 GB for `groHMM`, and 8.8 s and ca. 1 GB for the `megadepth` baseline (median over the 32 sub-groups). On the genome-wide run, `fastder` processed one tissue sub-group in ca. 1.8 minutes (median 106 s) with maximum resident set of ca. 30 GB (median near 18 GB) (Table S3).

## Discussion

`fastder` implements fast expressed-region detection with bounded memory requirements at genome scale. Reference-agnostic detection of expressed region from RNA-seq coverage, as provided by tools such as `derfinder` and `TAR-scRNA-seq` (grohMM) [13], has broadened the analysis of expression data beyond the reference annotation constraints, allowing the detection and quantification of noncanonical expression features, such as cryptic exons or alternate UTRs. `fastder`’s innovation rely on its increased computational performance, the native ingestion of `recount3` data, and the refining of expressed regions into multi-exon units by parsing the spliced reads (as provided by junction files) with, potentially, strand information from junction files.

`fastder`’s exon-level accuracy is competitive with coverage-only callers, such as `derfinder` and the `megadepth` baseline. `derfinder` additionally fits a statistical model for differentially expressed regions, which `fastder` does not. `groHMM` is sensitive at the base level but does not place exon boundaries on splice sites, being less competitive than `fastder` or `derfinder` in calling exon boundaries precisely. Performance on recovering transcripts is very low for `fastder`, so is for `derfinder` and `groHMM`, given their inability to reconstruct isoforms (a task they were not designed to address). `fastder` also calls multi-exon expressed regions and aims to assign strand information, if crossing junctions, extending coverage-only capabilities with splicing-specific ones.

As a coverage-based expressed region caller, `fastder` has several limitations. The requirement of pre-specifying a hard cut-off for gene expression to distinguish signal from background favors calling regions from highly expressed genes. Similarly, it neglects that different genes have different exon expression levels. The reliance on `STAR`-called junctions, both annotated and novel, makes it sensitive to `STAR` alignment tuning, including the two-pass mapping.

`fastder`’s user interface is a single command-line. As inputs, it requires fastqs or `recount3` identifiers. Optional flags set the chromosomes or scaffolds to use, the coverage and region length thresholds, and the number of threads to spawn. A planned next step is to run `fastder` across a large slice of `recount3` and publish the expressed-region catalog itself as a separate resource.

## Conclusion

`fastder` detects expressed regions from coverage bigWigs and `STAR` junction files as shipped by `recount3`. Optionally, it can be run from raw reads. As outputs, it produces a GTF with the called unstranded single exon and stranded multi-exon regions. Being 15-20x faster than `derfinder` and `groHMM`, `fastder` has a reduced memory fingerprint due to the use of sparse data representations, allowing the calling of expressed regions at scale.

## Availability and requirements

- **Project name:** `fastder`
- **Project home page:** https://github.com/imallona/fastder.
- **Operating system(s):** Platform independent
- **Programming language:** C++ (C++20)
- **Other requirements:** CMake 4.0 or newer and a C++20 compiler; optionally libBigWig with zlib and libcurl for direct bigWig input; otherwise `bigWigToBedGraph` from kentUtils.
- **License:** GPLv3
- **Any restrictions to use by non-academics:** none beyond the GPLv3 terms.

## Declarations

### Ethics approval and consent to participate

Not applicable. The study used public, de-identified RNA-seq data only.

### Consent for publication

Not applicable.

### Availability of data and materials

The `fastder` source code is at https://github.com/imallona/ `fastder` under GPLv3. The Snakemake evaluation workflow is at https://github.com/imallona/ `fastder_analysis`. The TDP-43 data are `recount3` study SRP166282 (GEO GSE121569). The GTEx data were served by `recount3`. Simulated data were generated with `ASimulatoR` and are packaged in the evaluation workflow.

### Authors’ contributions

ML wrote the original `fastder` implementation. TK wrote the original evaluation workflow. IM extended the tool and the evaluation pipeline, and drafted the manuscript. All authors read and approved the final manuscript.

### Usage of LLMs

The authors declare having used Microsoft Copilot for code reviewing and grammar checks.

## Acknowledgements.

We thank Mark D. Robinson for access to compute and feedback.

## Supplementary methods

### Inputs and data model

`fastder` consumes four file types (Fig. S1A) from the `recount3` resource or from the `recount3`-like workflow (provided with the tool). The coverage files are per-sample bigWig files, a binary format storing continuous RNA-seq coverage along the genome, that lack strand information.

Splice junctions are supplied as exon-exon junction counts in Matrix Market format, an MM file and an RR file. The MM file is a sparse matrix with one line per (junction, sample) pair carrying the junction count. The RR file has one line per junction, with coordinates, strand, splice motif and an annotation flag; its line number is the junction identifier used by the MM file.

For `recount3`-reliant runs, `fastder` reuses `recount3`’s sample identifiers, including: the `sample_id` of the MM file, an internal `rail_id`, and a per-source `external_id`. A metadata CSV carries these identifiers, the study name, and the URL of each coverage file. `fastder` reads this CSV to map between the identifiers and to find which junctions were called in the samples the user supplied.

### Parsing and concurrency

During the parsing stage the small RR and metadata CSV files are read with `std::getline` and formatted extraction. The MM file, which can hold about 700 million lines of three tab-separated integers for `recount3` runs, is read line by line and converted with `std::from_chars`. The bedGraph resulting from the read of the coverage bigWig via Kent utils [12] is read so the chromosome column is extracted by pointer, and its three integer columns are converted with `std::from_chars`.

The Parser class stores the user-specified chromosomes and the supplied sample identifiers `unordered_set` containers, providing constant-time lookup. Per-base coverage and MM data are stored in `unordered_map` keyed by chromosome, which allows parallelization by chromosome later. Coverage parsing is multi-threaded: one thread parses one bedGraph and moves to the next free file as soon as it is done, so no thread stays idle while files remain. An atomic index, safe to share across threads, assigns each chromosome to exactly one thread for the mean coverage and bump hunting stages. For a full human genome run, `fastder` is best suited with about 23 cores, one per autosomal or X chromosome.

### Simulations

To simulate expression with known splicing properties, we ran `ASimulatoR` with five samples per scenario and on human chr21 only. We simulated fastqs at 5, 10, 30 and 40 million reads depth. The four `fastder` parameters and their defaults were: `min_coverage` 0.05 CPM, the coverage threshold; `min_length` 10 bp, the minimum expressed region length; `position_tolerance` 5 bp, the allowed offset between an expressed region edge and a junction coordinate; and `coverage_tolerance`, which gates stitching on coverage similarity and is left effectively off by default, so stitching is driven solely by junctions. The benchmark explored the `min_coverage`, `min_length` and `position_tolerance` values in an expanded grid.

### gffcompare and fuzzy performance metrics

To evaluate methods performance using the reference simulations as truths, we used `gffcompare` metrics, including precision and sensitivity at the base, exon, intron, intron-chain, transcript and locus levels, and custom fuzzy metrics, more lenient on exact boundaries. The fuzzy metrics included: the best exonic Jaccard overlap of each true transcript against any call on the same strand; the signed base-pair distance from each called exon boundary to the nearest true boundary; the locus recall defined as the fraction of true loci with at least a given fraction of their exonic length covered; and strand concordance, being either concordant, discordant, unstranded or unmatched to the truth.

## Supplementary figures

**Figure S1:**
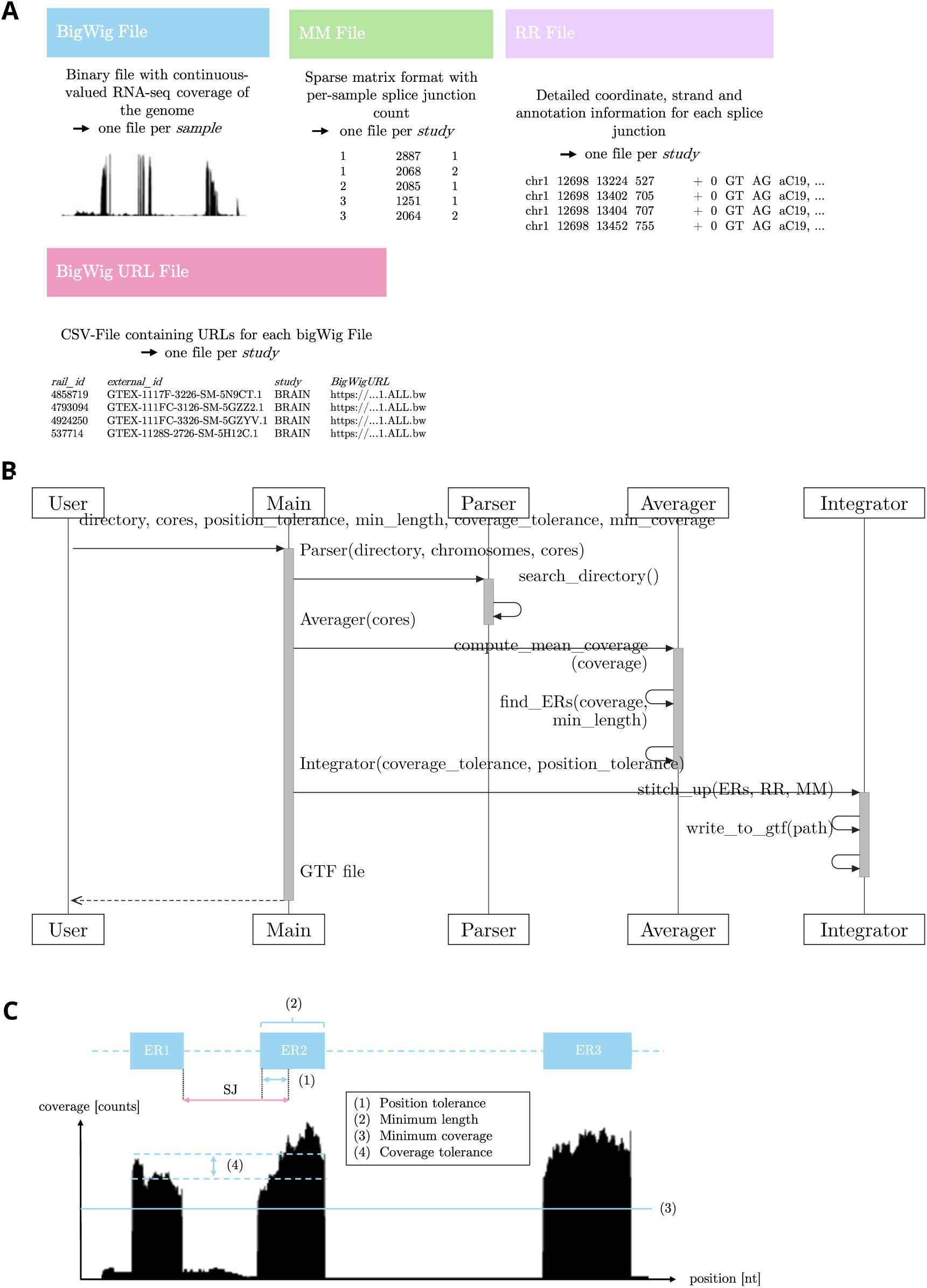
Overview of the `fastder` software. **(A)** The four input file types provided to `fastder`: bigWig, MM and RR file (splice junctions) and bigWig URL CSV file. **(B)** Sequence diagram illustrating the three main functional stages of `fastder`. The diagram begins with the user input, continues with file parsing, averaging, bump hunting and, finally, the integration of splice junction files. `fastder` writes the results to a GTF file. **(C)** Visualization of the parameters *position tolerance, minimum length, minimum coverage, coverage tolerance*. These parameters define what qualifies as an expressed region in terms of required coverage, length, and offset of expressed region (ER) and splice junction (SJ) coordinates.

**Figure S2:**
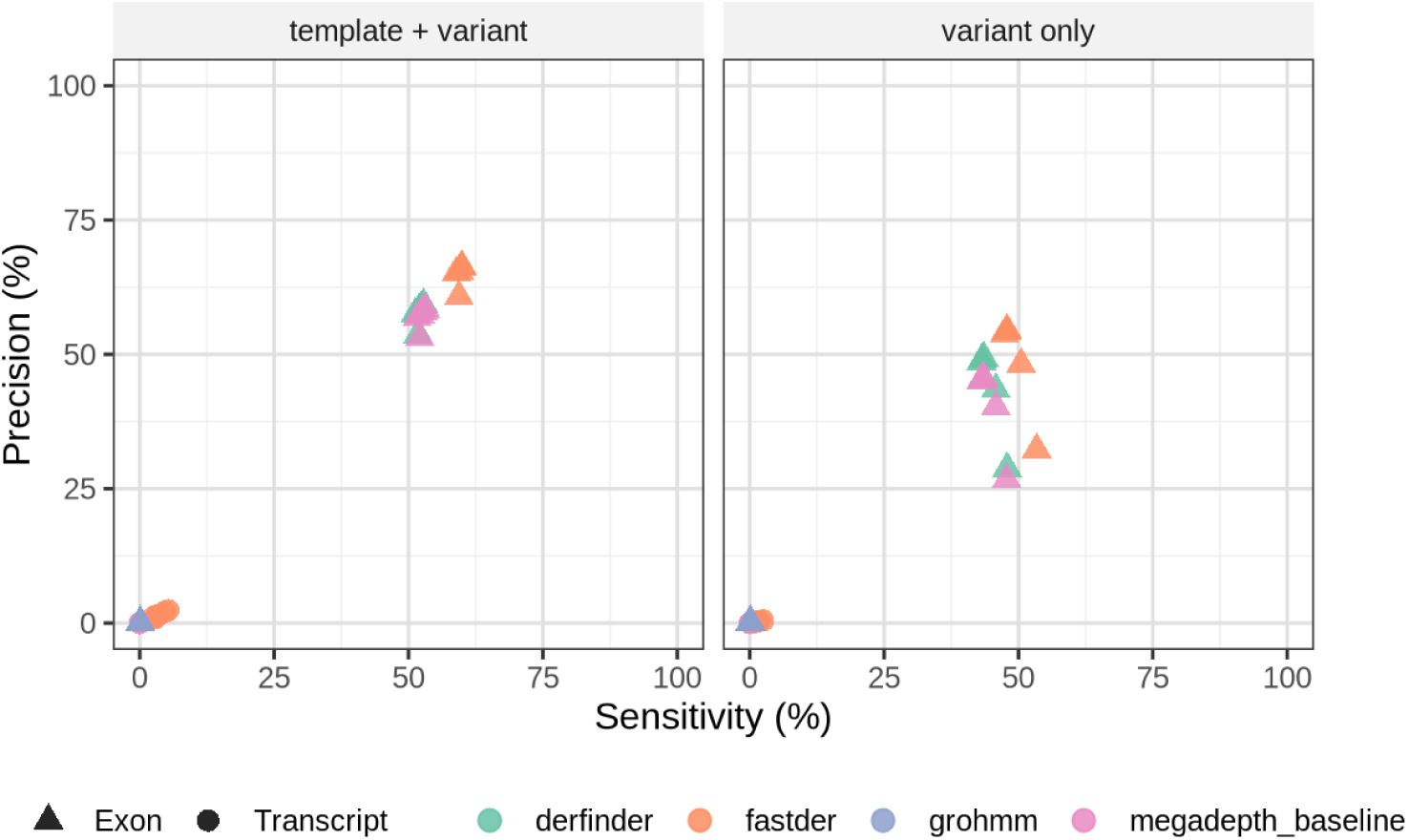
Precision and sensitivity in simulated data. Each point is one tool and one alternative splicing event sample, at the exon and transcript levels of `gffcompare`. Up and to the right is better.

**Figure S3:**
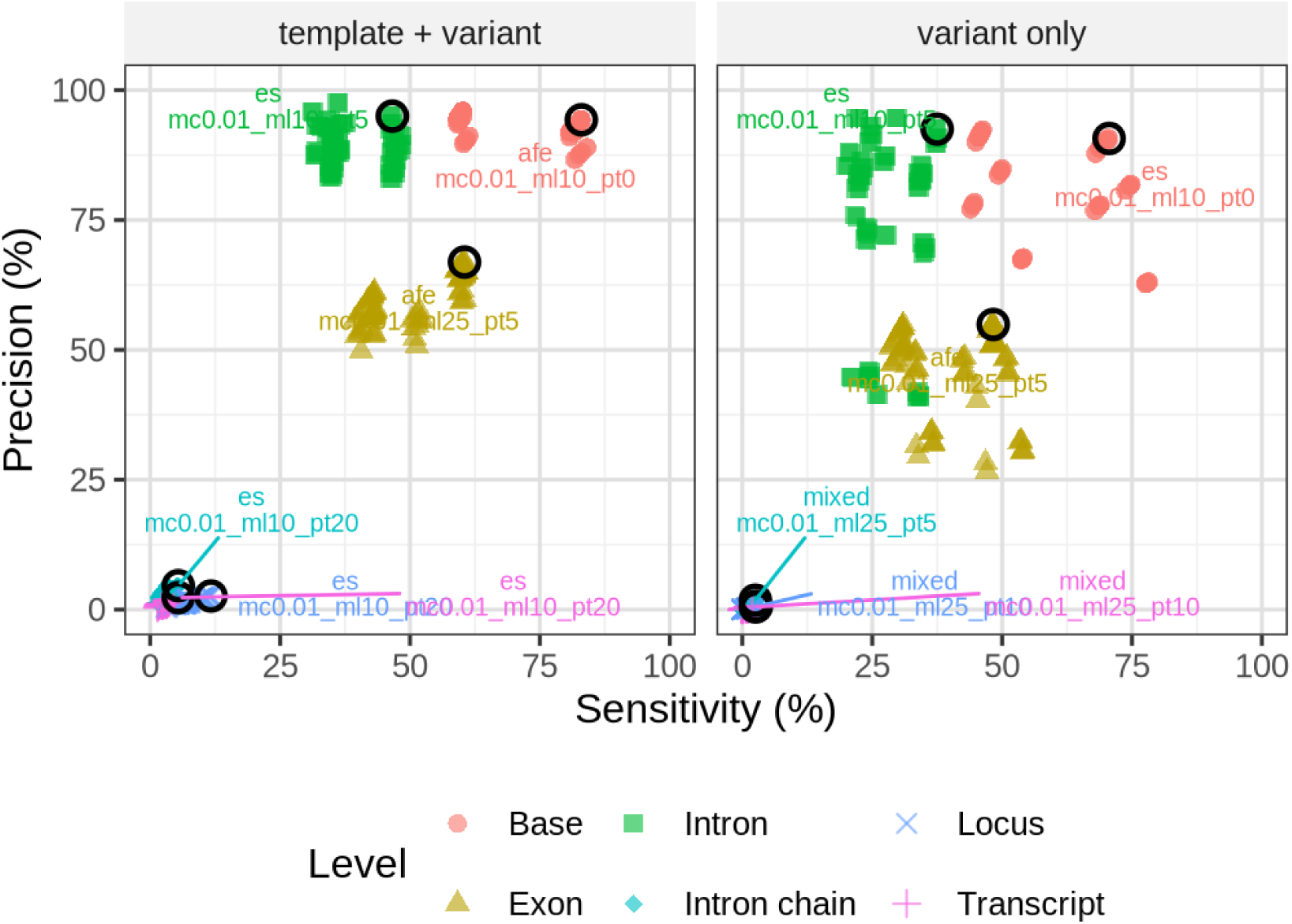
Precision and sensitivity in simulated data (`fastder` only) across the `gffcompare` base, exon, intron, intron chain, locus and transcript level, split by scenario. Top performing hyperparameters are circled and labelled. Up to the right is better.

**Figure S4:**
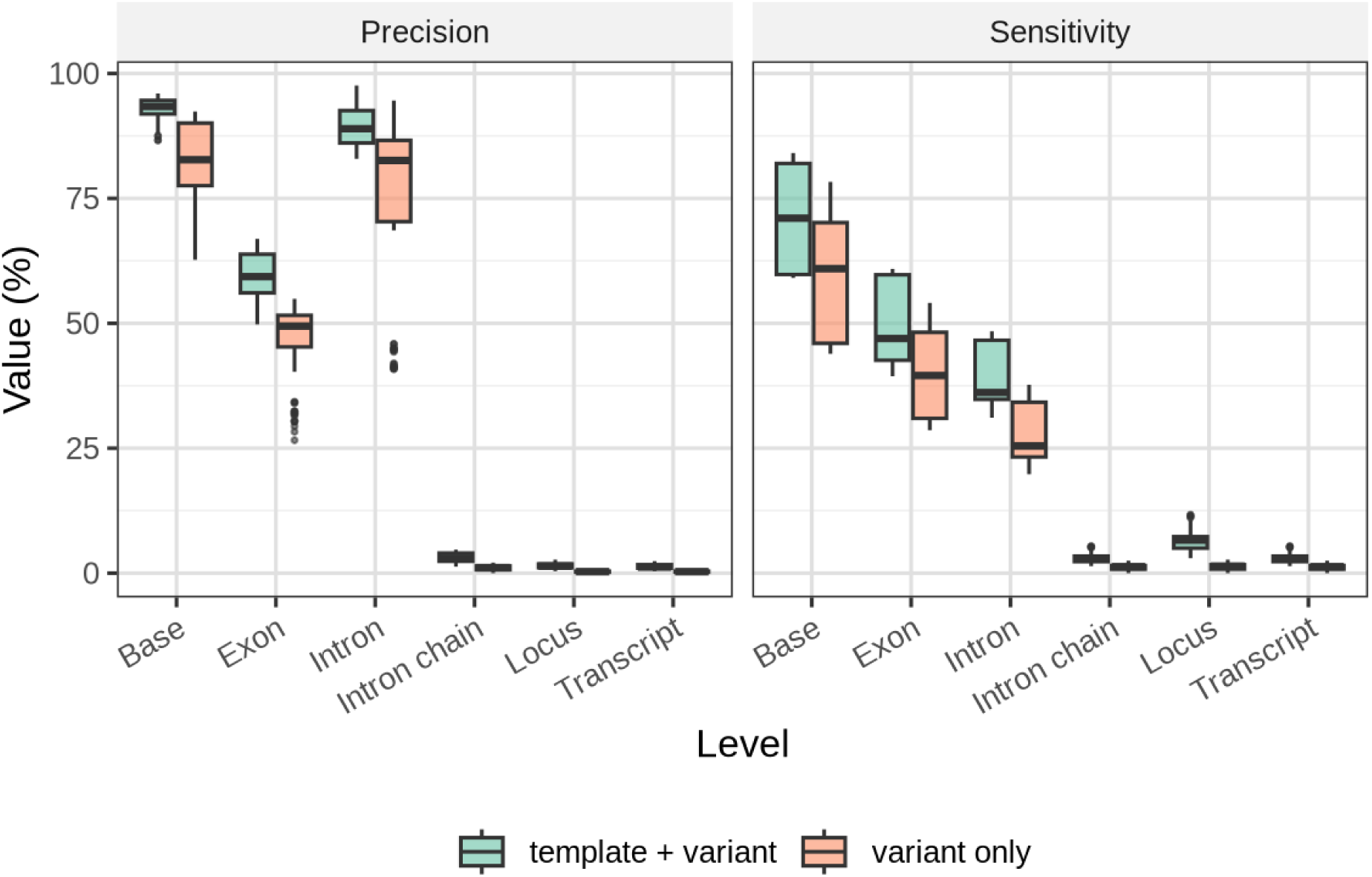
Per-level sensitivity and precision bars. The transcript, intron-chain and locus levels carry non-zero values only for `fastder`, because the coverage-only baselines emit single-exon regions and `groHMM` segments do not snap to exon edges.

**Figure S5:**
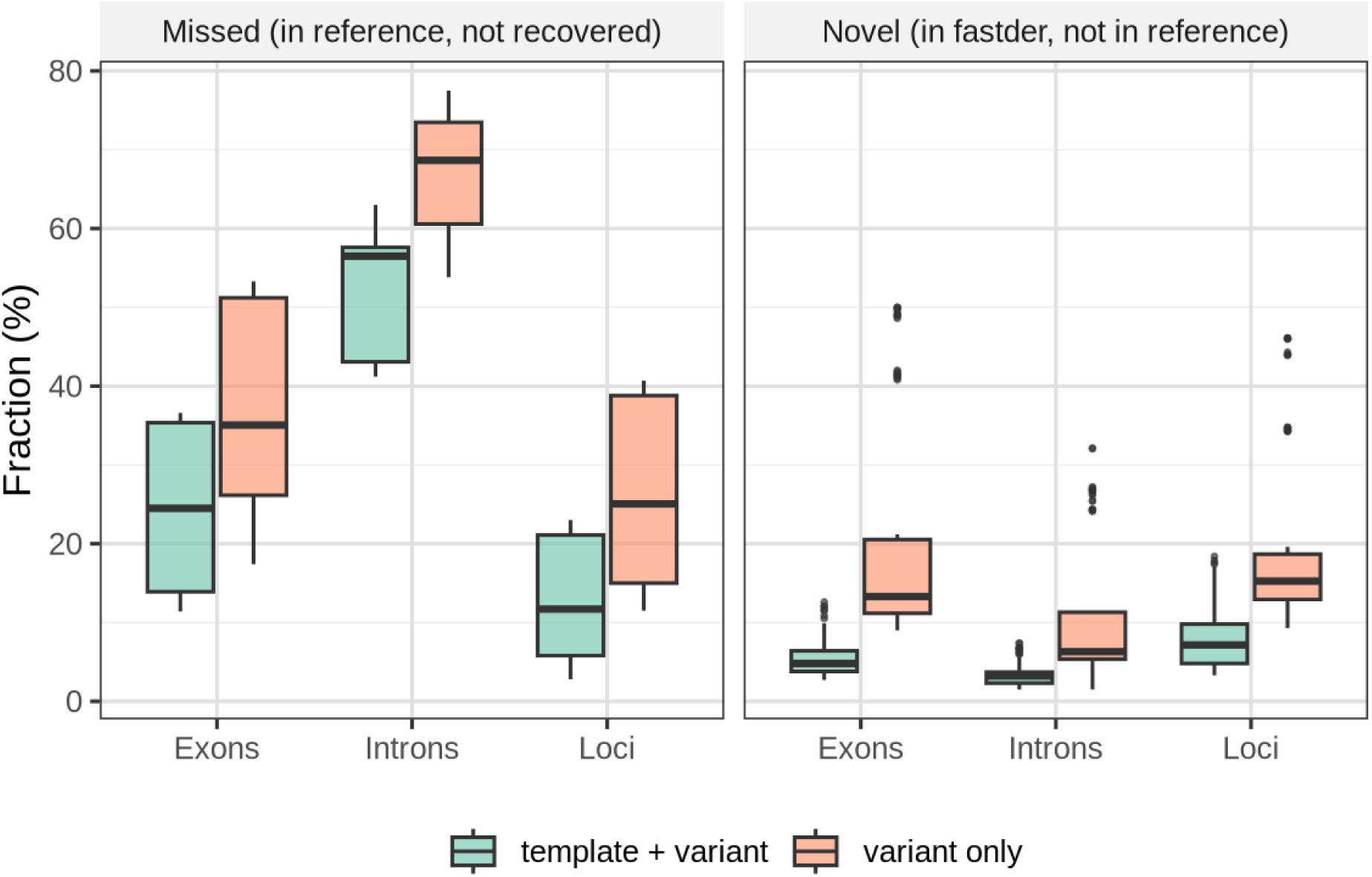
Missed and novel features per tool under the two simulation regimes (reference and variant; and variant only). Missed features are false negatives, novel features are false positives.

**Figure S6:**
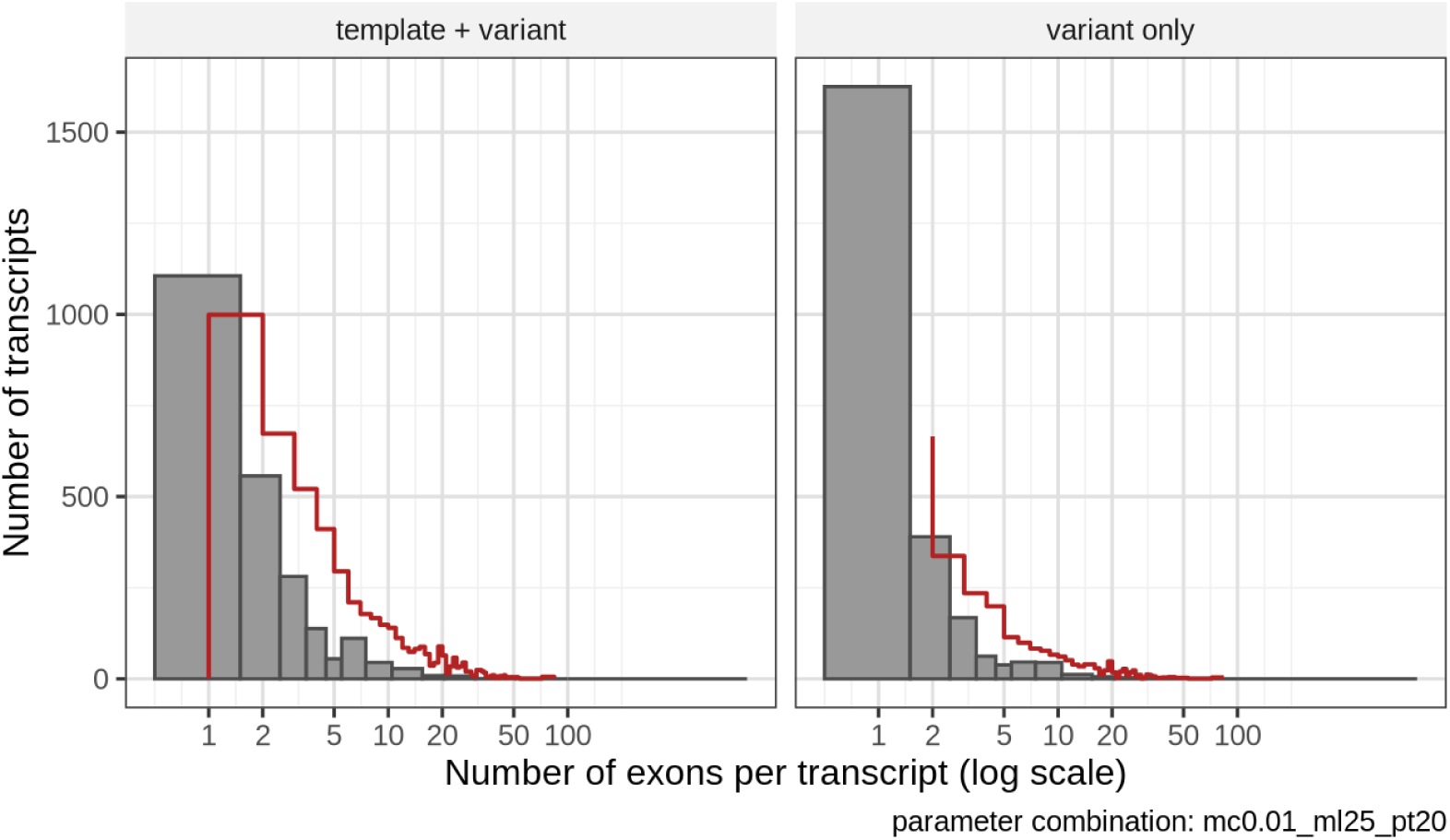
Number of exons per expressed region as called by `fastder`. Most regions are monoexonic.

**Figure S7:**
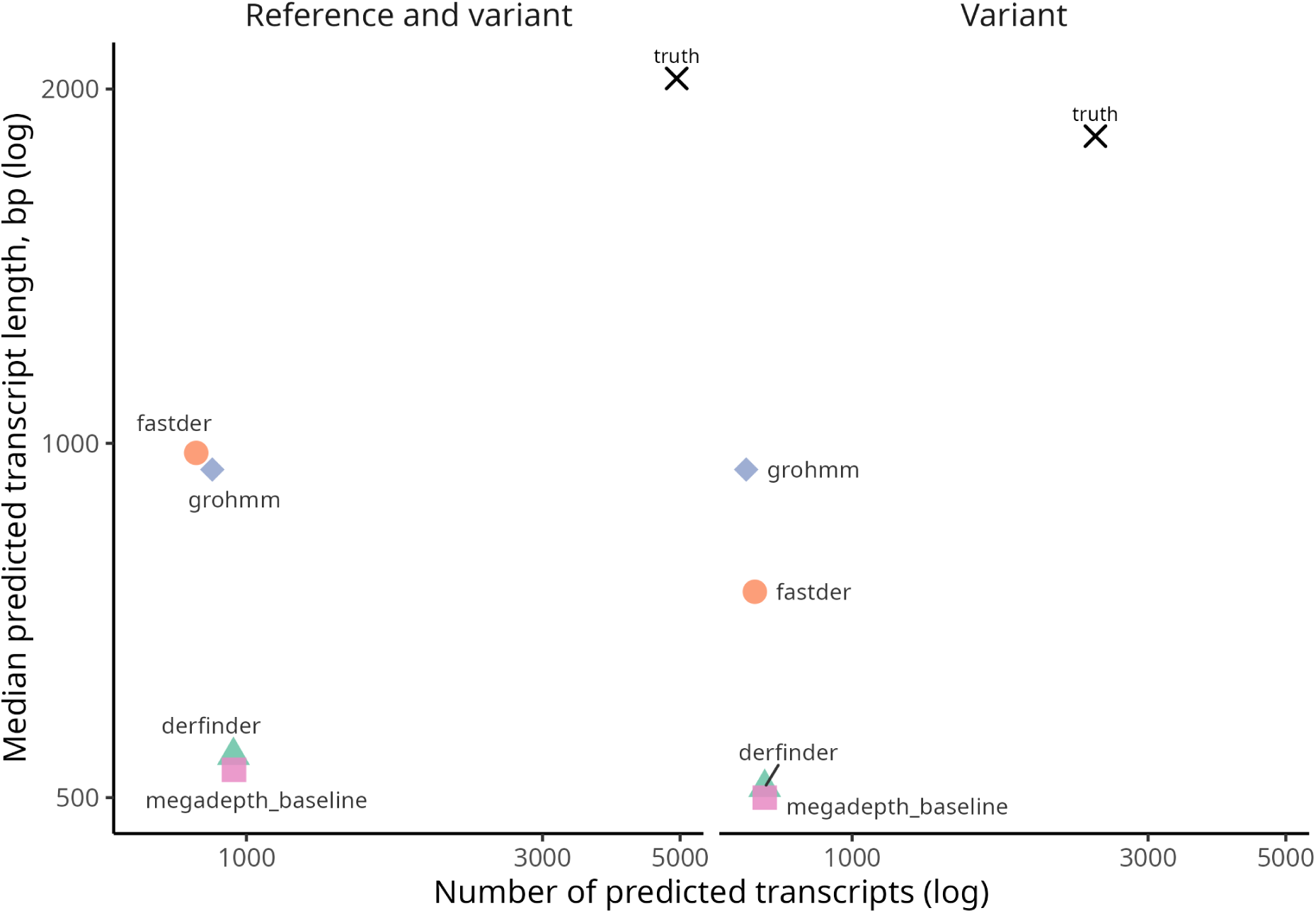
Expressed regions granularity. Dots depict each tool’s performance in terms of counts and length of the called expressed regions. The true (simulated) number is highlighted with a cross.

**Figure S8:**
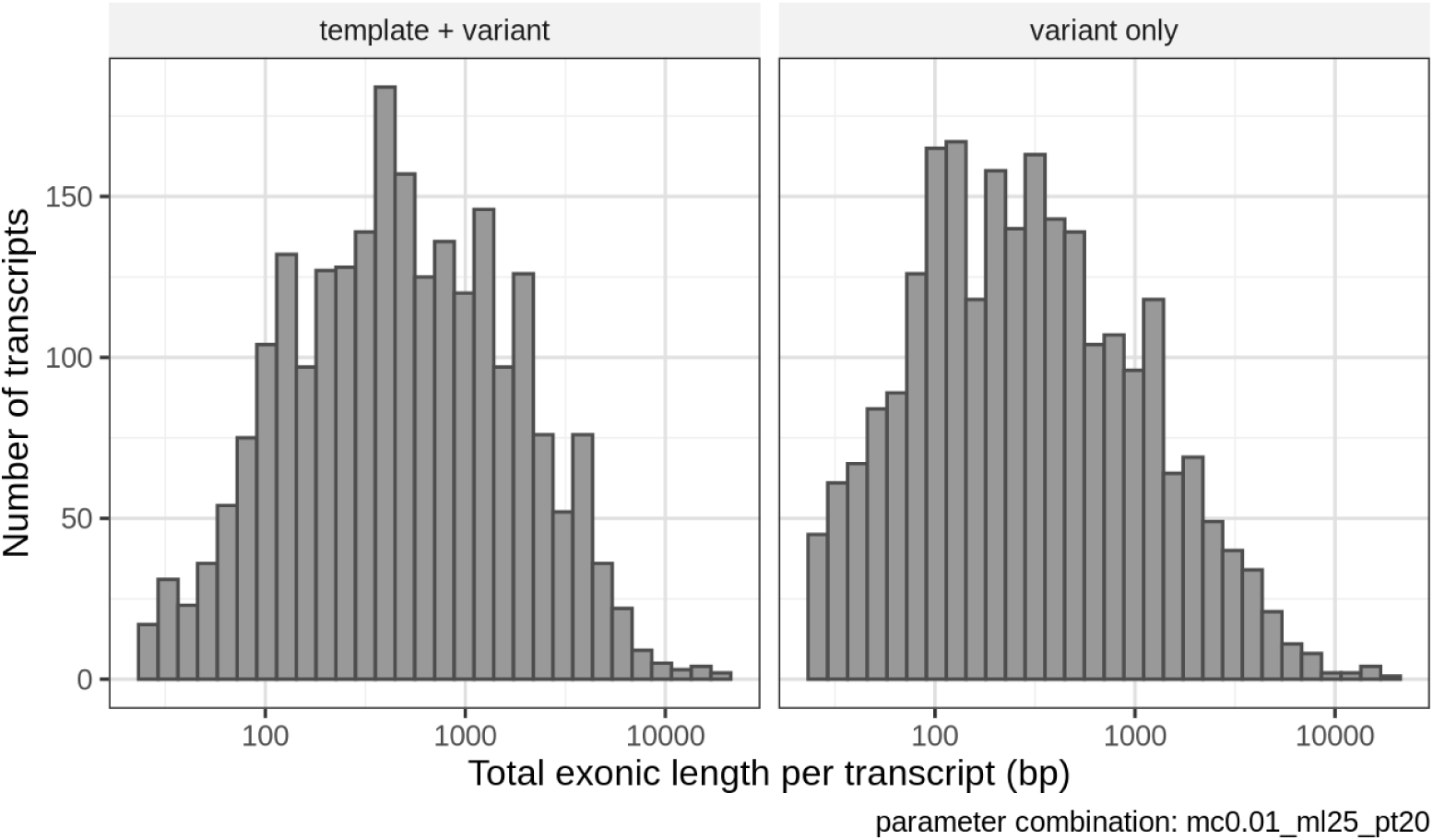
Exonic length per `fastder` region.

**Figure S9:**
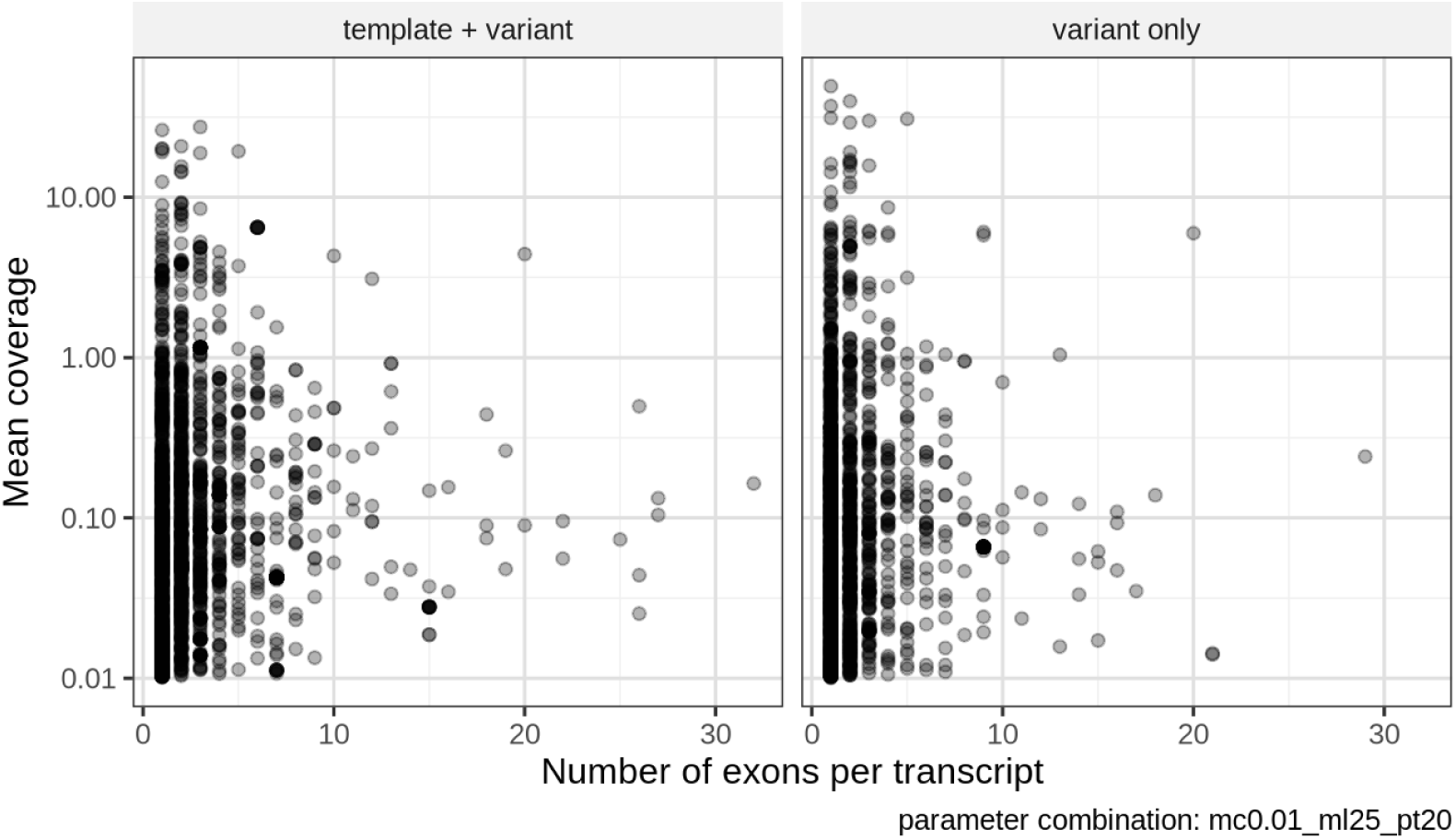
Mean coverage against the number of exons, per called `fastder` region.

**Figure S10:**
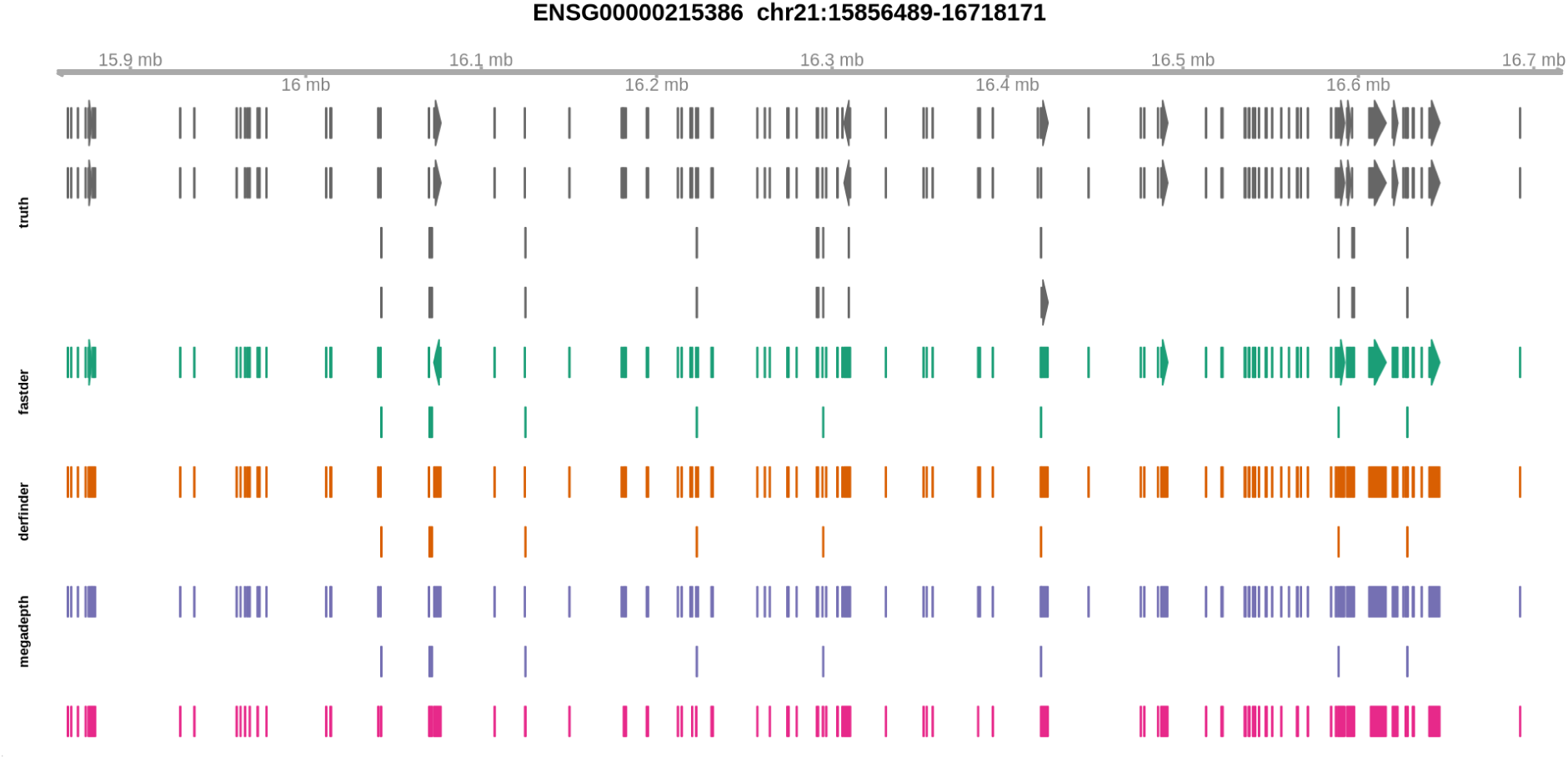
Genome track of a true multi-exon expressed region and each tool’s calls. Blocks that line up with the truth have accurate boundaries.

**Figure S11:**
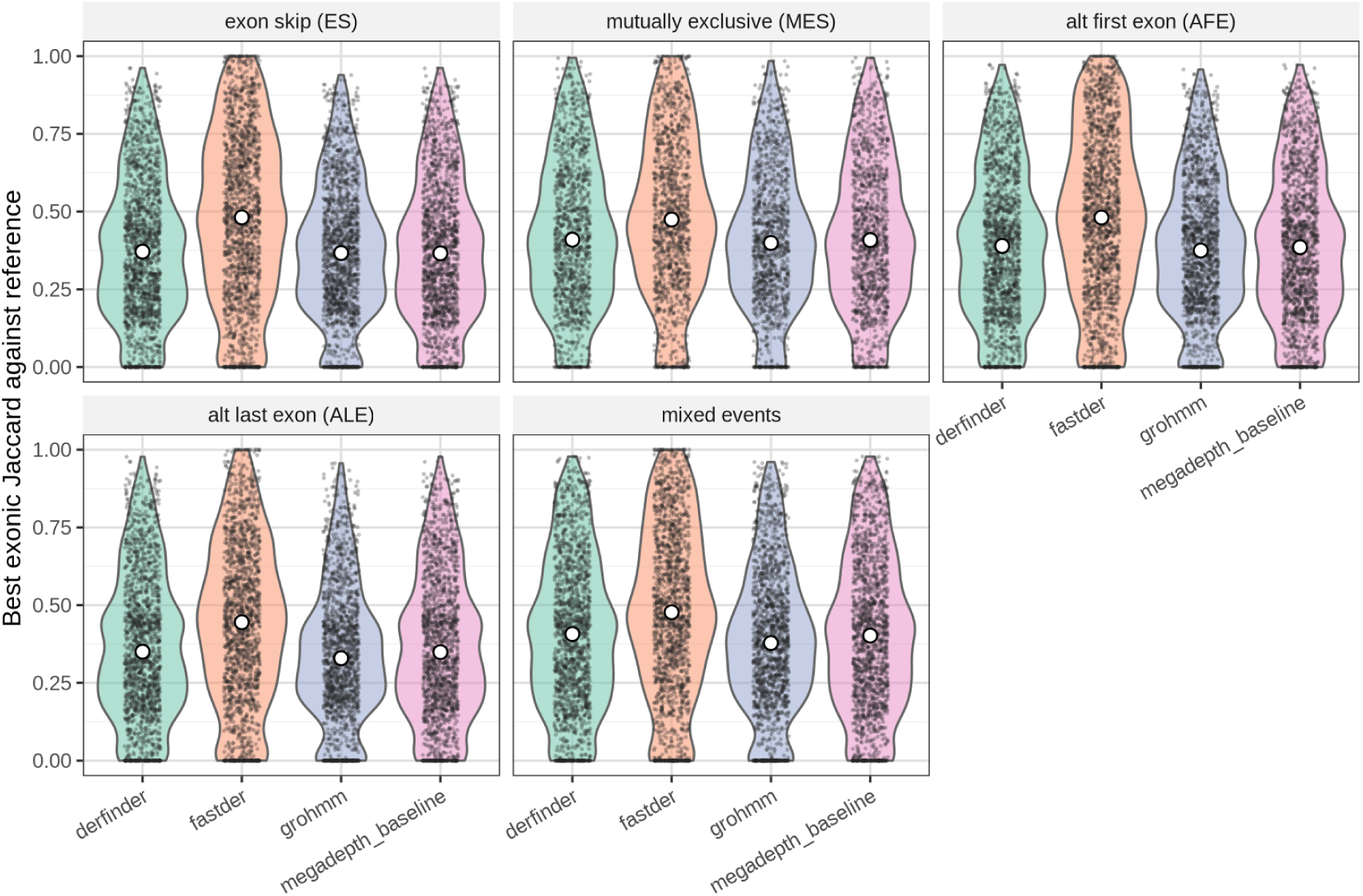
Best exonic Jaccard overlap per tool, by alternative splicing event sample and simulation scenario.

**Figure S12:**
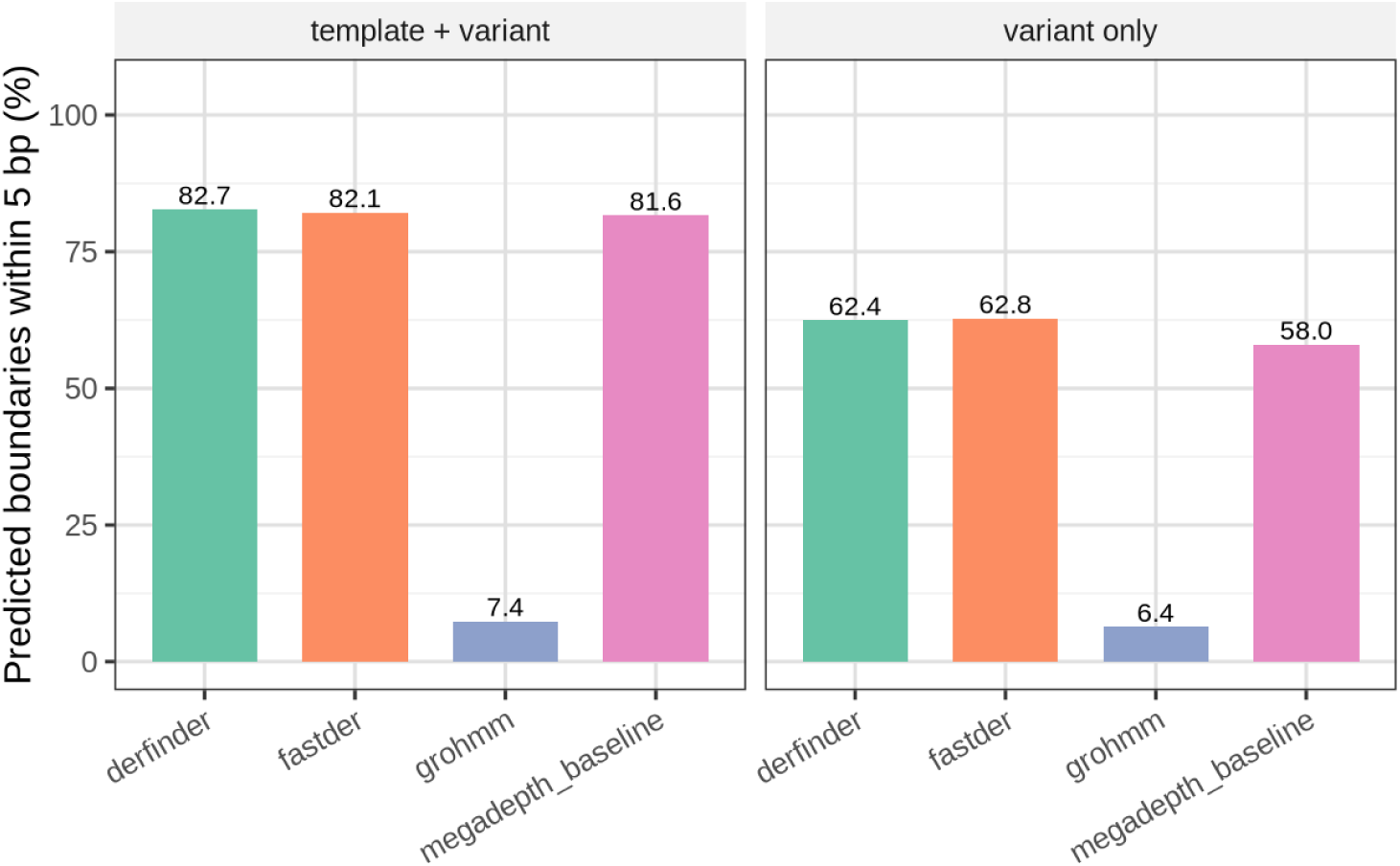
Boundary precision at 5 bp. Fraction of predicted exon boundaries within 5 bp of a true splice site, per tool and simulation scenario.

**Figure S13:**
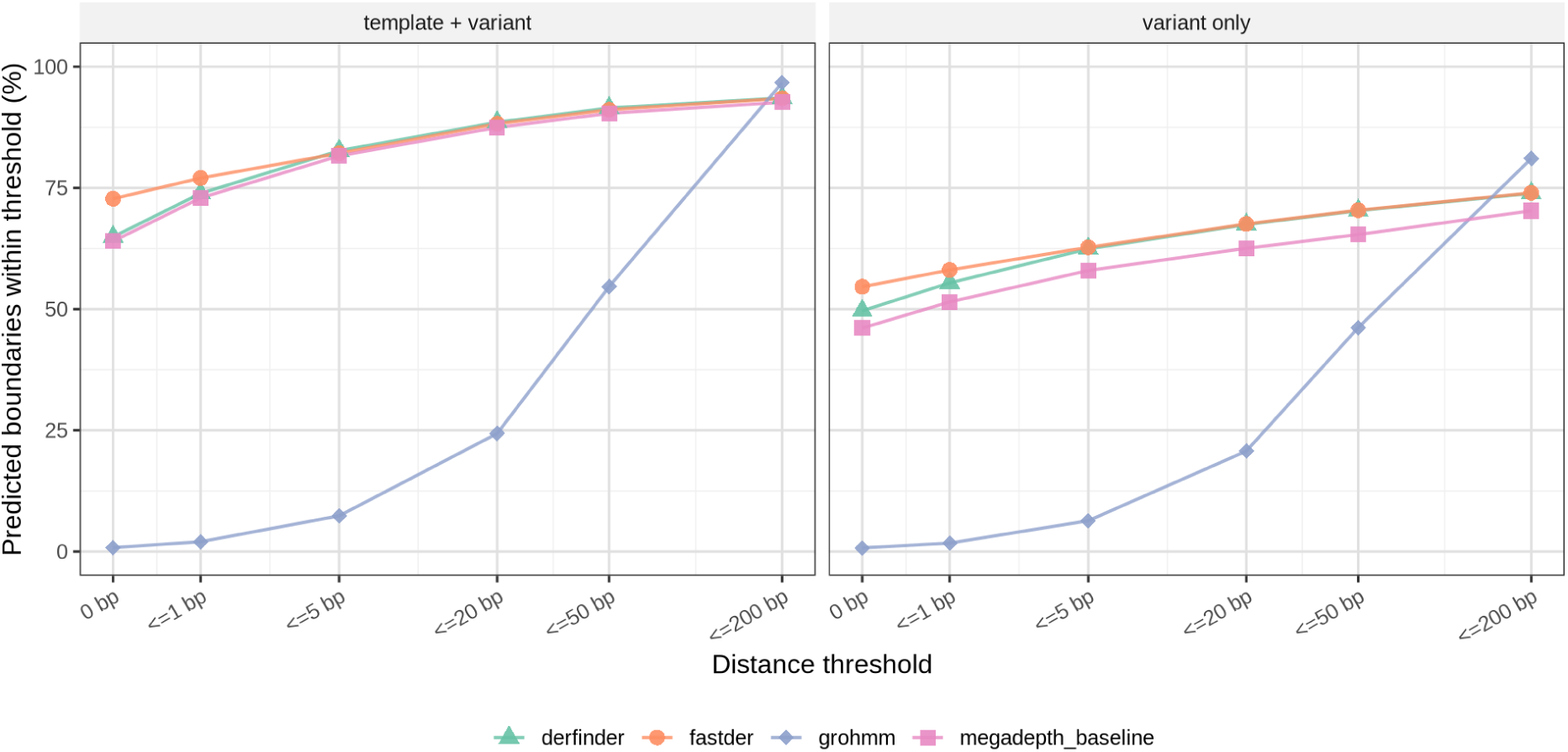
Boundary precision (fraction of predicted exon boundaries) at different distance tolerances to the true splice site. Facetted by variant-only and template plus variant scenarios.

**Figure S14:**
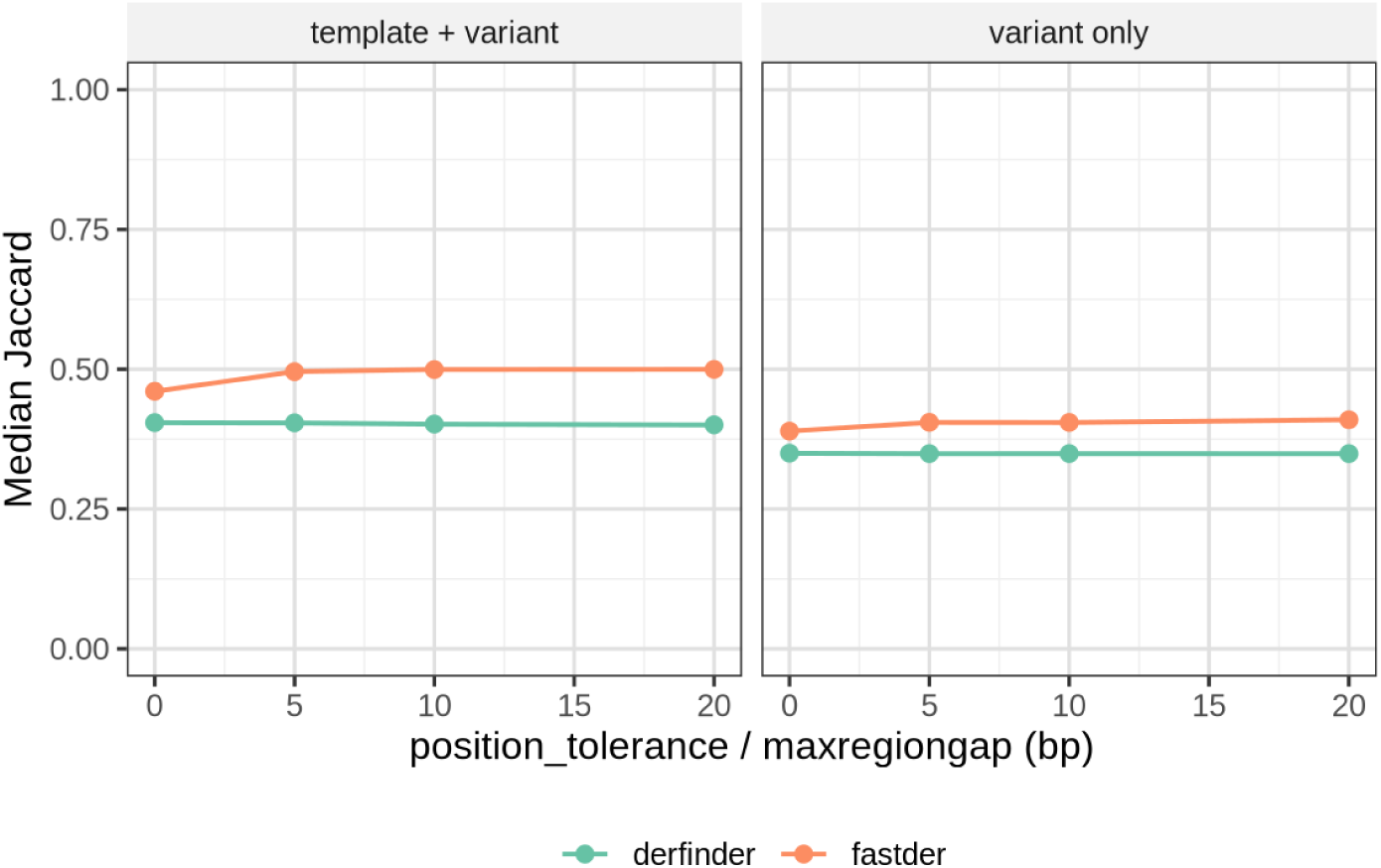
Effect of the boundary-distance tolerance hyperparameters on the fuzzy match rate, per tool. Exploring `fastder`’s `position_tolerance` and `derfinder`’s `maxregiongap` from 0 to 20 bp do not impact the median exonic Jaccard overlap against the simulated truth.

**Figure S15:**
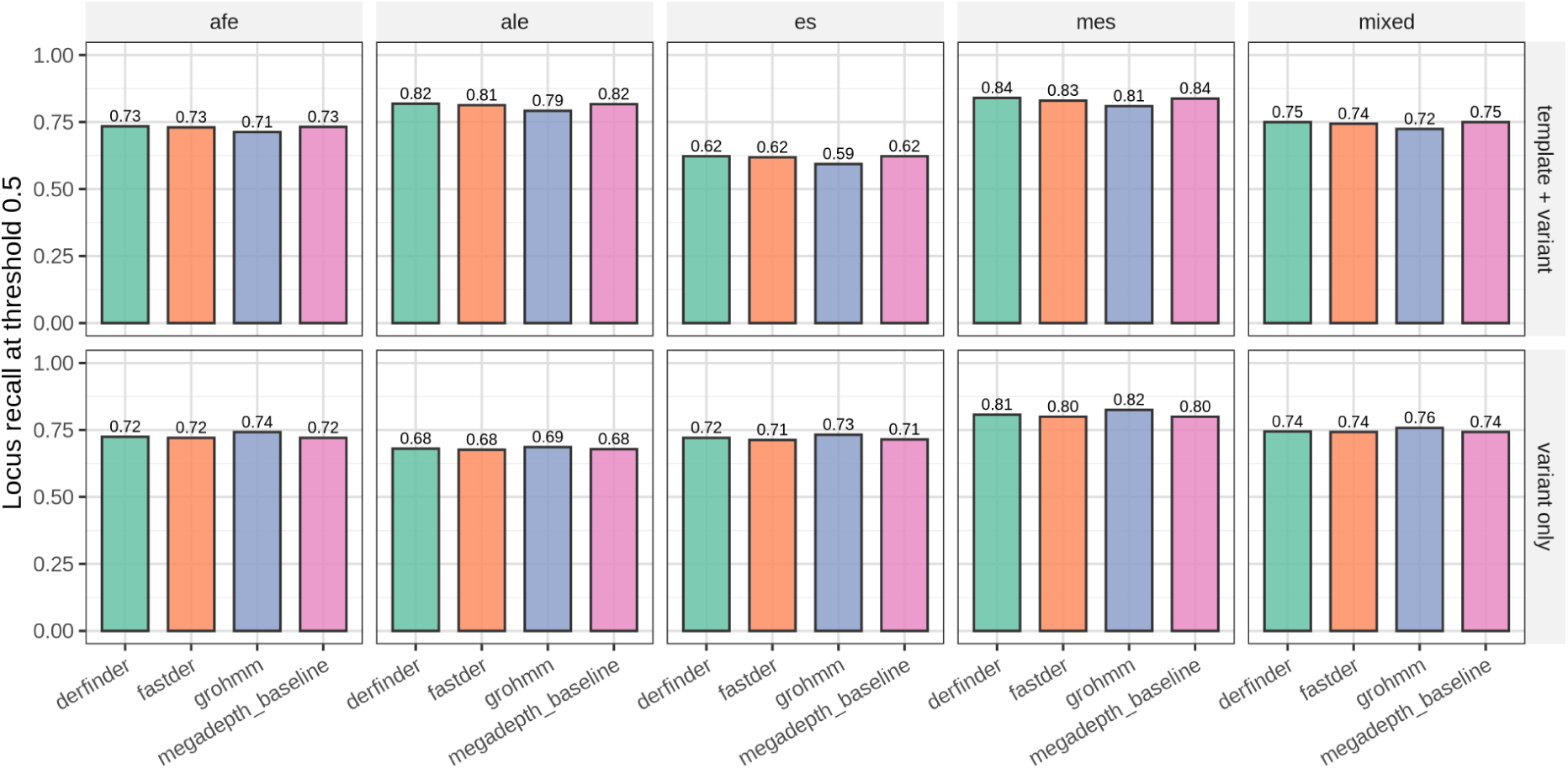
Locus recall (fraction of true loci with at least half of their exonic length covered by calls) at an exonic-overlap threshold of 0.5, by alternative splicing event and simulation scenario.

**Figure S16:**
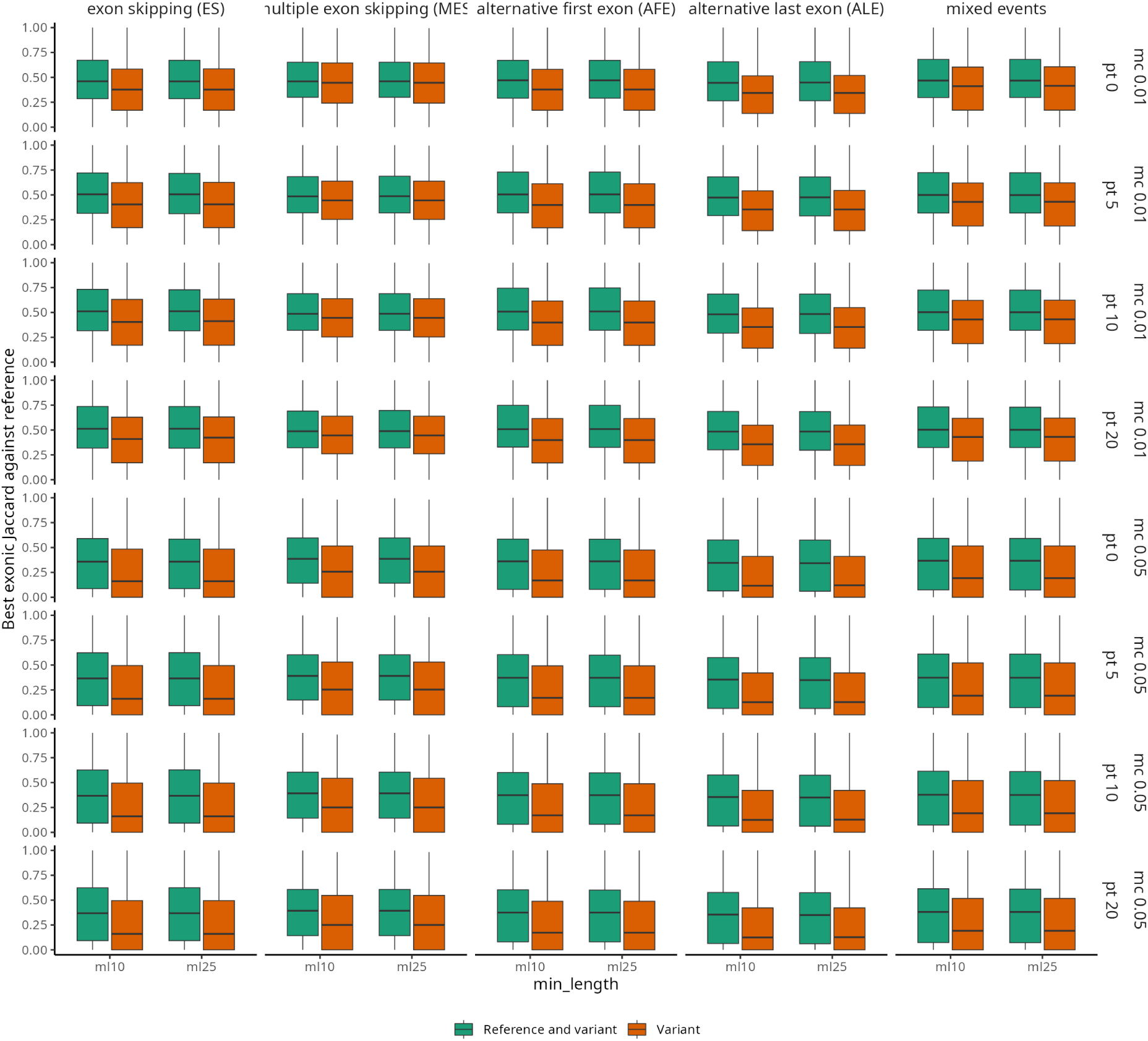
Per-event exonic Jaccard overlap of `fastder` calls against the simulated truth set, decomposed over the parameter grid. The panel columns are the AS event class and the rows are `min_coverage`; within each panel the boxes run over `min_length` and `position_tolerance`, split by scenario.

**Figure S17:**
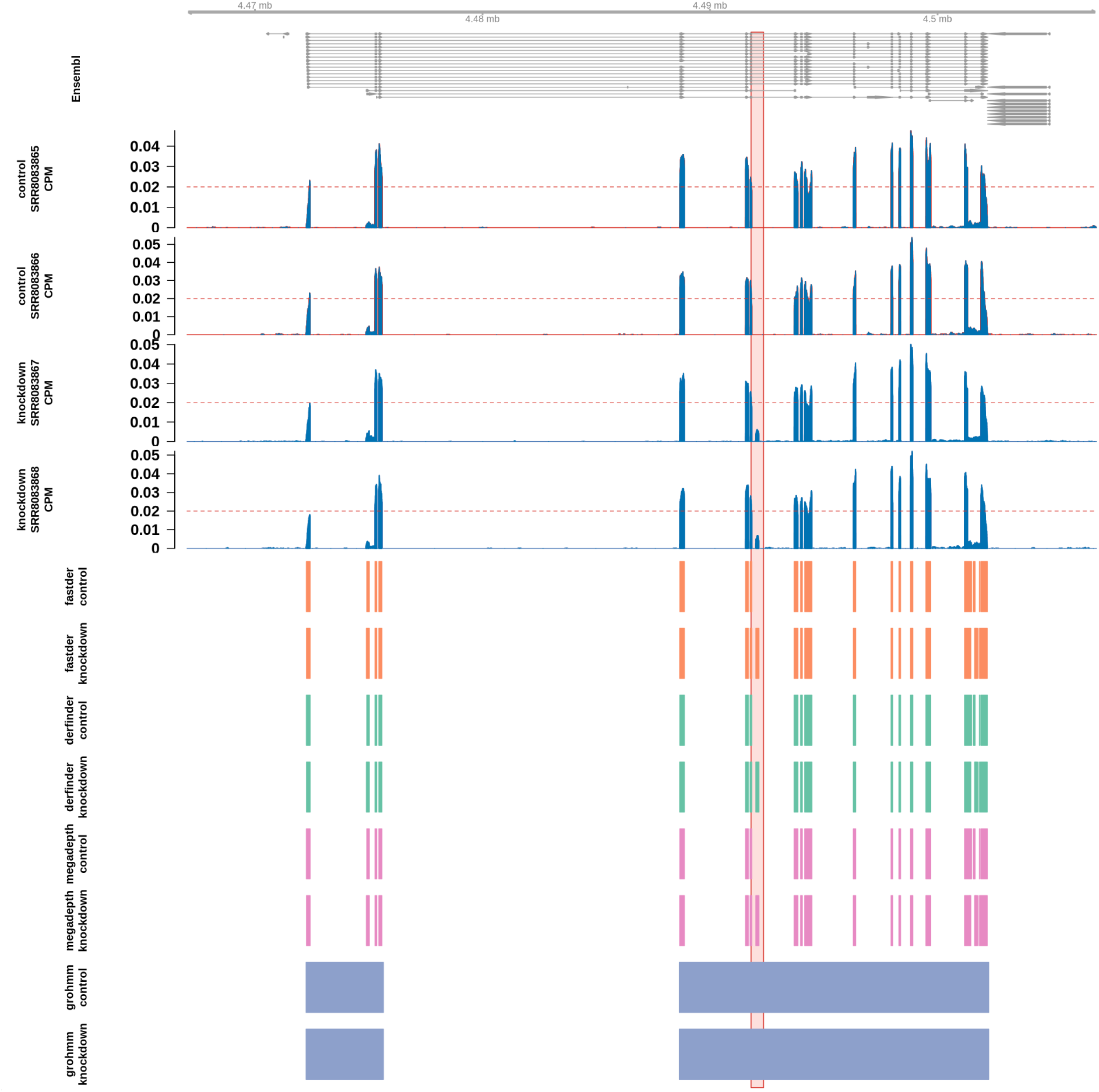
HDGFL2 cryptic exon (box) as detected across tools.

**Figure S18:**
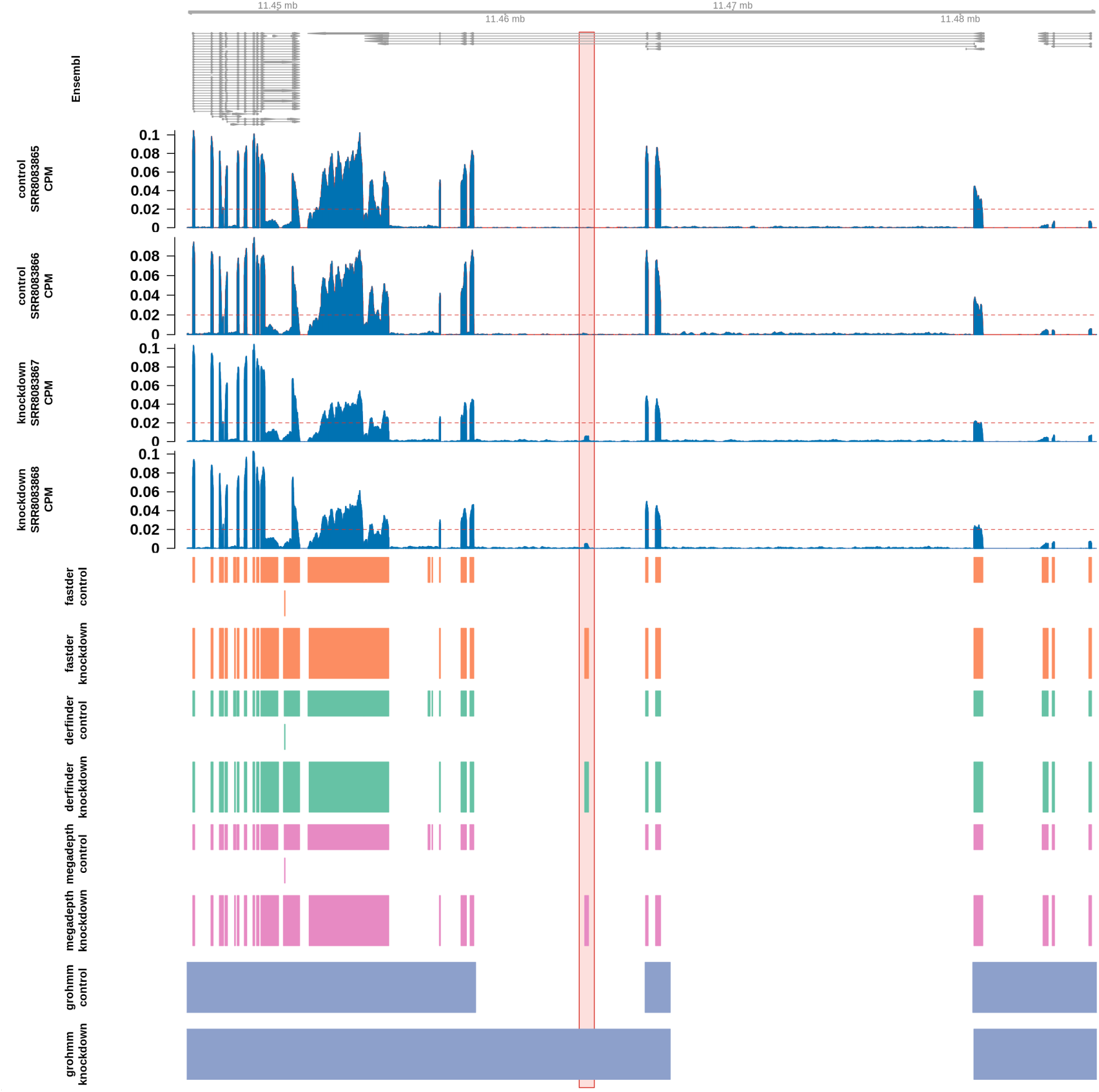
ELAVL3 cryptic exon (box) as detected across tools.

**Figure S19:**
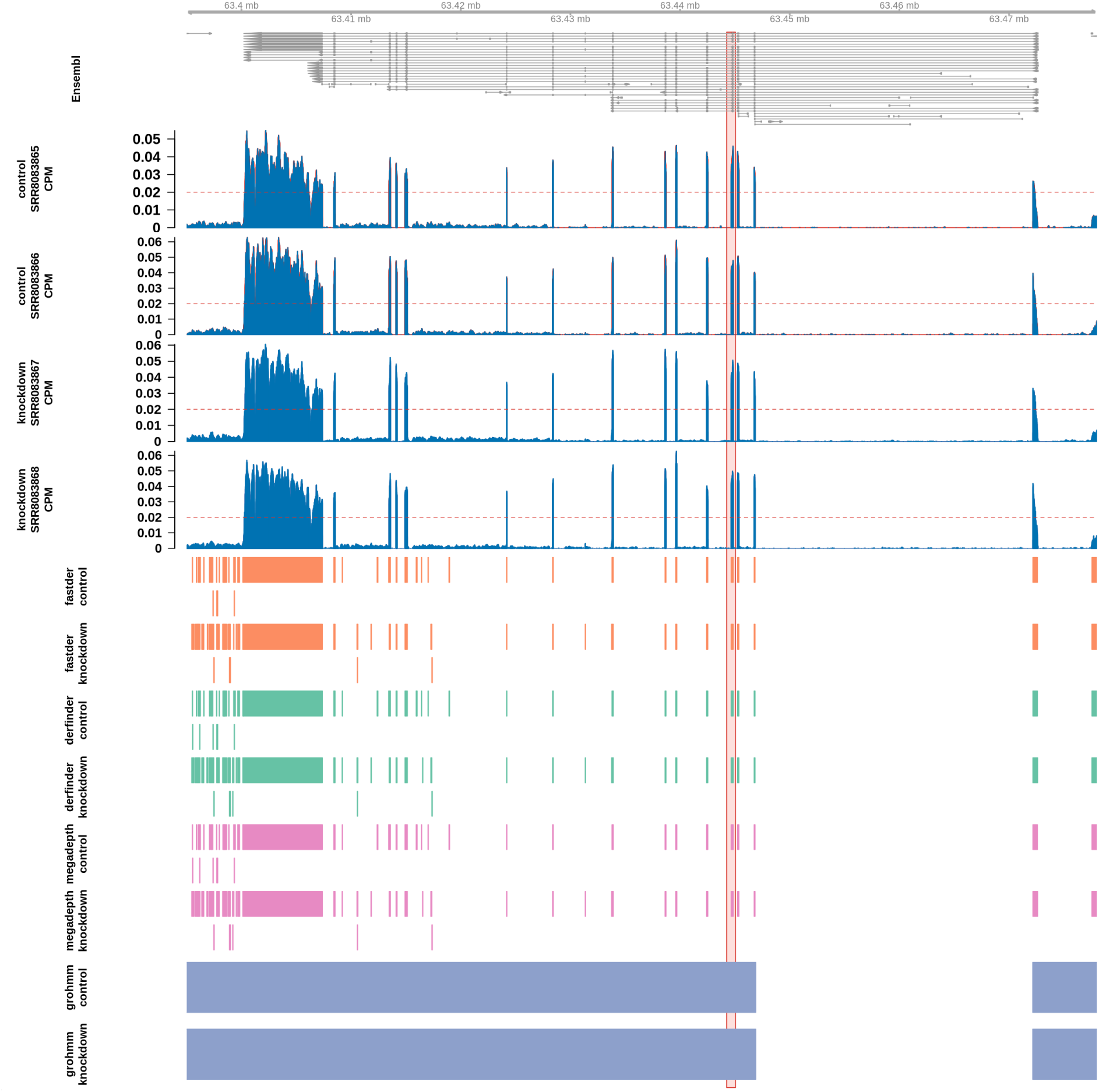
KCNQ2 cryptic exon (box) as detected across tools.

**Figure S20:**
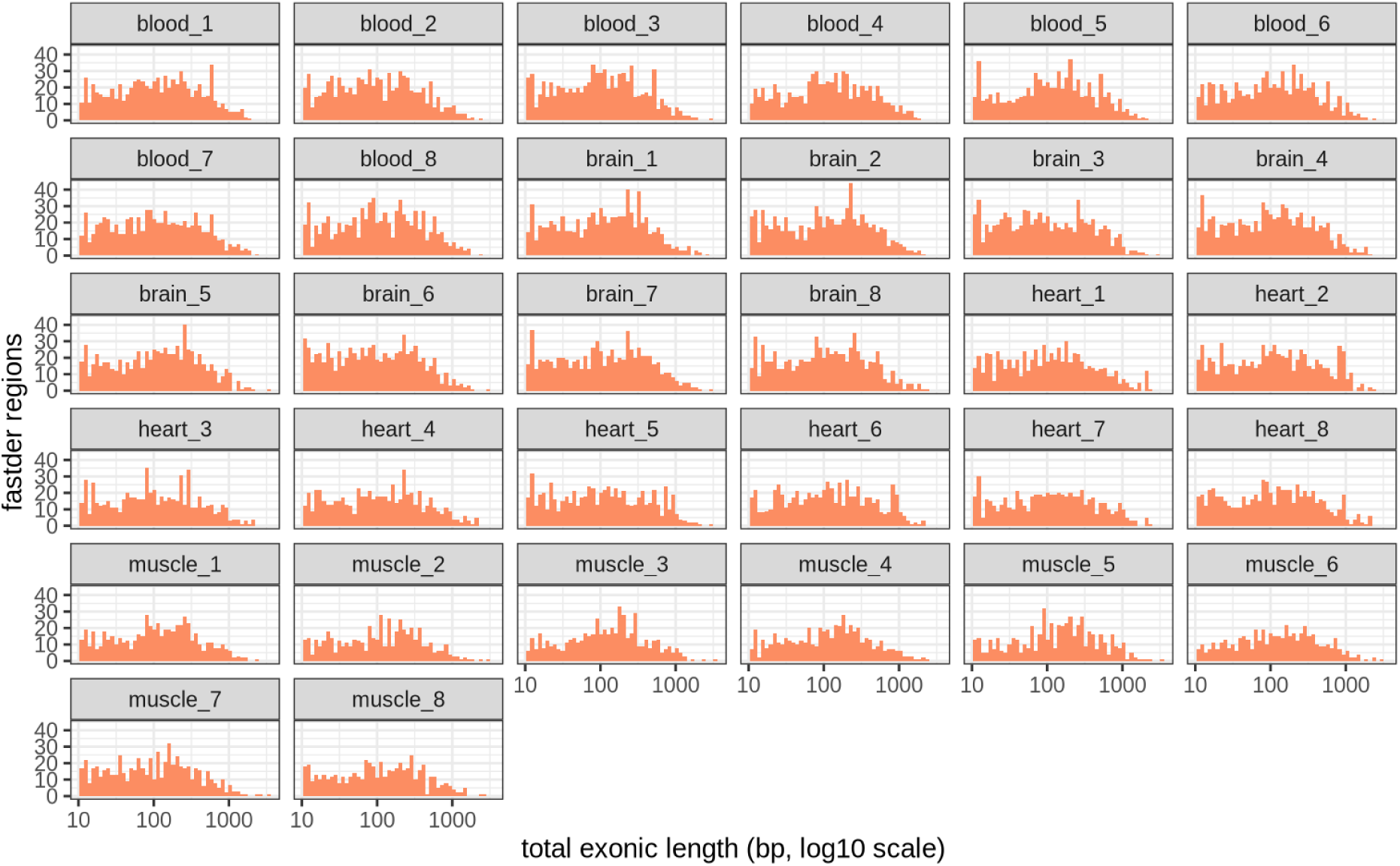
Exonic length per `fastder` region on the GTEx dataset.

**Figure S21:**
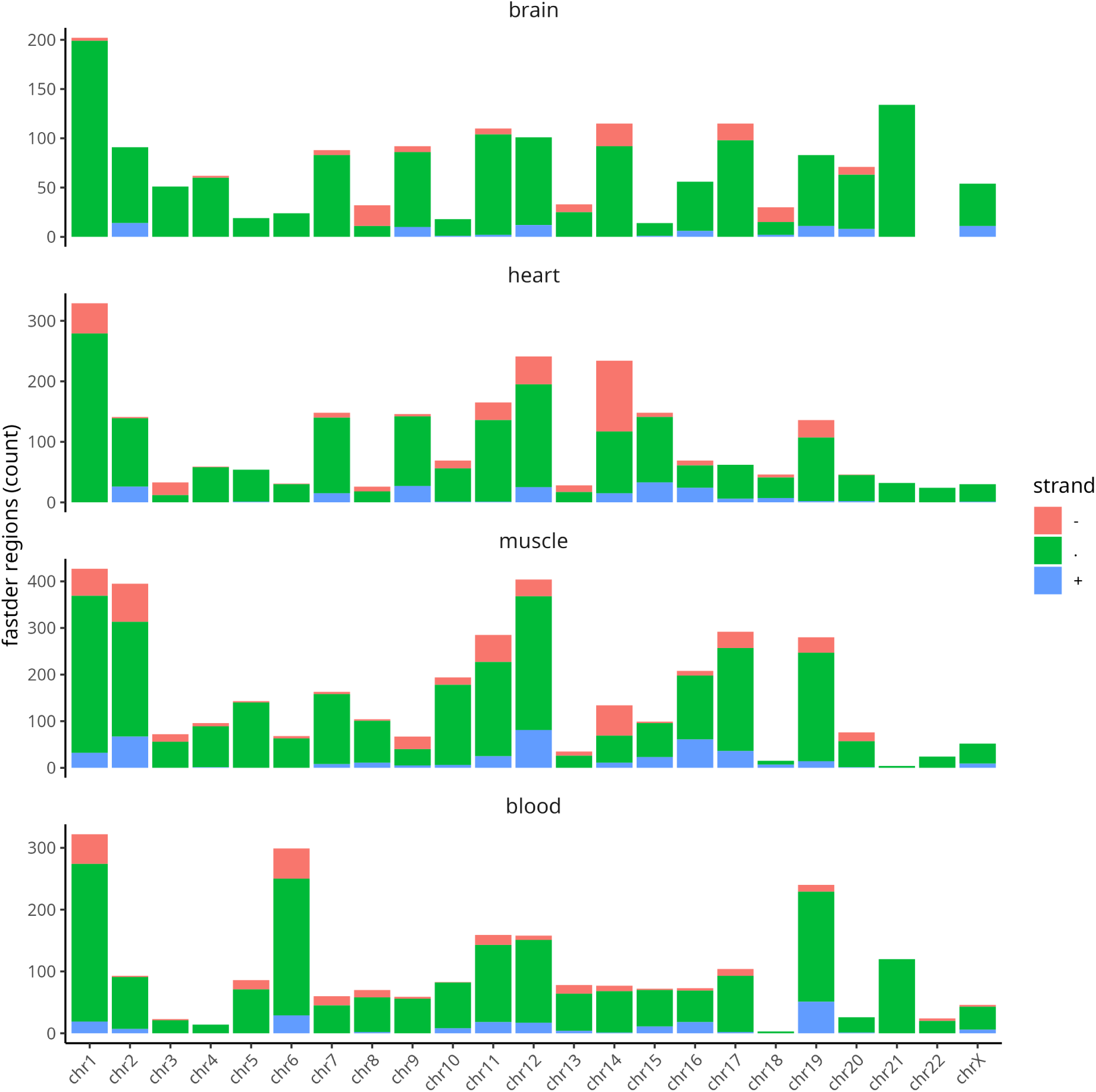
Genomic distribution of `fastder` regions on the GTEx tool-comparison run, by chromosome, tissue, and strand. Coverage threshold was kept at 1 CPM (hence only calling highly expressed regions).

## Supplementary tables

**Table S1:**
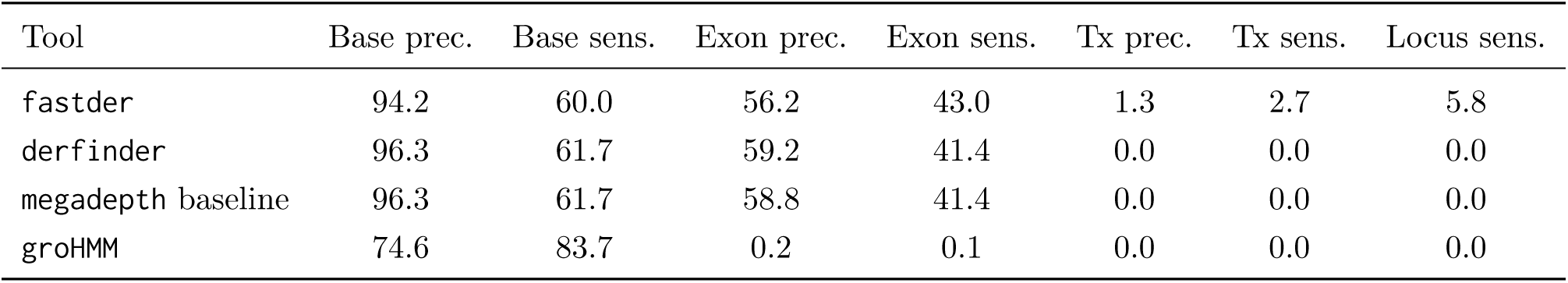
Precision and sensitivity on simulated RNA-seq (`ASimulatoR`, chr21, 10 million reads per sample, template-and-variant scenario), per tool at default parameters (`min_coverage` 0.05 CPM, `min_length` 10 bp, `position_tolerance` 5 bp), mean over five alternative splicing event samples. `fastder` is shown with junction stitching enabled. Prec.: precision (percent); sens.: sensitivity (percent).

**Table S2:** Exon precision across the sequencing-depth sweep (`ASimulatoR`, chr21, template-and-variant scenario, default parameters, mean over the five AS event samples). Values are exon precision in percent, plus `fastder` exon sensitivity. `groHMM` is omitted because its exon precision stays near zero at every depth (its HMM segments do not snap to exon boundaries). Prec.: precision; sens.: sensitivity.

| Depth | fastder exon prec. | derfinder | megadepth baseline | fastder exon sens. |
| --- | --- | --- | --- | --- |
| 5M | 53.7 | 52.7 | 50.3 | 43.1 |
| 10M | 56.2 | 59.2 | 58.8 | 43.0 |
| 30M | 61.5 | 59.1 | 58.1 | 43.5 |
| 40M | 62.1 | 59.2 | 58.8 | 43.5 |

**Table S3:** Wall-clock time and peak resident memory per run and tool, median, min and max over the benchmarked samples. Simulation rows are at default cross-tool parameters; GTEx chr19 is the four-tool comparison run, GTEx genome-wide and the two TDP-43 runs are `fastder` only. Peak RSS is the maximum resident set reported by the Snakemake `benchmark` directives. * No reliable peak RSS: for quick runs, the execution completes faster than benchmark data pooling.

| Run | Tool | Wall clock (s) | Peak RSS (GB) |
| --- | --- | --- | --- |
| Simulation 5M | **fastder** | 2.2 [1.9–2.6] | 0.6 [0.3–0.8] |
|  | **derfinder** | 60.2 [55.2–65.2] | 2.9 [2.8–3.1] |
|  | **groHMM** | 70.1 [63.3–79.8] | 1.5 [1.1–1.5] |
|  | **megadepth baseline** | 6.0 [5.6–6.3] | 0.7 [0.7–0.8] |
| Simulation 10M | **fastder** | 2.7 [2.2–3.2] | 0.8 [0.7–0.9] |
|  | **derfinder** | 60.0 [53.6–66.4] | 2.6 [2.4–2.8] |
|  | **groHMM** | 68.0 [61.8–76.2] | 1.5 [1.0–1.5] |
|  | **megadepth baseline** | 6.3 [5.8–6.8] | 0.8 [0.7–0.9] |
| Simulation 30M | **fastder** | 3.2 [2.5–3.9] | NA* |
|  | **derfinder** | 69.3 [59.7–78.8] | 2.8 [2.8–2.8] |
|  | **groHMM** | 72.4 [61.8–83.3] | 1.2 [1.1–1.5] |
|  | **megadepth baseline** | 7.8 [7.6–8.1] | 0.8 [0.6–1.0] |
| Simulation 40M | **fastder** | 3.3 [2.8–3.8] | 1.3 [1.3–1.3] |
|  | **derfinder** | 71.9 [61.4–82.4] | 2.9 [2.8–3.0] |
|  | **groHMM** | 71.2 [62.8–82.8] | 1.2 [1.2–1.5] |
|  | **megadepth baseline** | 7.6 [6.8–8.3] | 0.8 [0.6–0.9] |
| GTEx chr19 | **fastder** | 4.1 [3.0–5.6] | 1.3 [0.4–2.0] |
|  | **derfinder** | 57.5 [49.1–88.1] | 3.3 [1.3–3.8] |
|  | **groHMM** | 75.6 [73.4–89.5] | 1.3 [1.1–1.7] |
|  | **megadepth baseline** | 8.7 [7.2–16.0] | 1.2 [0.8–1.6] |
| GTEx genome-wide | **fastder** | 74.9 [46.7–101.7] | 15.5 [8.0–30.0] |
| TDP-43 (STMN2) | **fastder** | 4.7 [4.2–5.3] | 1.7 [1.5–2.0] |
| TDP-43 (panel) | **fastder** | 4.5 [4.1–4.9] | 1.7 [1.7–1.8] |

**Table S4:** `fastder` accuracy by `ASimulatoR` alternative-splicing event class (chr21, 10 million reads per sample, default parameters, junction stitching enabled), template-and-variant and variant-only scenarios. Prec.: precision (percent); sens.: sensitivity (percent).

| AS event class | Template and variant |  | Variant only |  |
| --- | --- | --- | --- | --- |
|  | Exon prec. | Exon sens. | Exon prec. | Exon sens. |
| Exon skipping (es) | 56.2 | 42.3 | 50.8 | 31.0 |
| Multiple exon skipping (mes) | 53.1 | 43.2 | 32.0 | 36.8 |
| Alternative first exon (afe) | 57.7 | 43.5 | 51.1 | 31.2 |
| Alternative last exon (ale) | 57.3 | 43.2 | 50.1 | 30.6 |
| Mixed | 56.9 | 42.9 | 46.2 | 33.7 |

**Table S5:** GTEx chr19 structural comparison against the Ensembl annotation (gffcompare, 32 per-sub-group catalogs, median). Prec.: precision (percent); sens.: sensitivity (percent).

| Tool | Base prec. | Exon prec. | Exon sens. | Tx prec. | Tx sens. |
| --- | --- | --- | --- | --- | --- |
| fastder | 98.8 | 51.8 | 2.1 | 5.2 | 0.1 |
| derfinder | 99.3 | 46.6 | 2.1 | 0.2 | 0.0 |
| megadepth baseline | 99.3 | 46.6 | 2.1 | 0.2 | 0.0 |
| groHMM | 33.9 | 1.7 | 0.0 | 2.7 | 0.1 |

